# MS^2^Rescore: Data-driven rescoring dramatically boosts immunopeptide identification rates

**DOI:** 10.1101/2021.11.02.466886

**Authors:** Arthur Declercq, Robbin Bouwmeester, Aurélie Hirschler, Christine Carapito, Sven Degroeve, Lennart Martens, Ralf Gabriels

**Affiliations:** VIB-UGent Center for Medical Biotechnology, VIB, Belgium; Department of Biomolecular Medicine, Ghent University, Belgium; Laboratoire de Spectrométrie de Masse BioOrganique (LSMBO), Université de Strasbourg, CNRS

**Keywords:** bioinformatics, immunopeptidomics, machine learning, proteomics, mass spectrometry, peptide identification

## Abstract

Immunopeptidomics aims to identify Major Histocompatibility Complex-presented peptides on every cell that can be used in anti-cancer vaccine development. However, existing immunopeptidomics data analysis pipelines suffer from the non-tryptic nature of immunopeptides, complicating their identification. Previously, peak intensity predictions by MS^2^PIP and retention time predictions by DeepLC, have been shown to improve tryptic peptide identifications when rescoring peptide-spectrum matches with Percolator. However, as MS^2^PIP was tailored towards tryptic peptides, we have here retrained MS^2^PIP to include non-tryptic peptides. Interestingly, the new models not only greatly improve predictions for immunopeptides, but also yield further improvements for tryptic peptides. We show that the integration of new MS^2^PIP models, DeepLC, and Percolator in one software package, MS^2^Rescore, increases spectrum identification rate and unique identified peptides with 46% and 36% compared to standard Percolator rescoring at 1% FDR. Moreover, MS^2^Rescore also outperforms the current state-of-the-art in immunopeptide-specific identification approaches. Integration of immunopeptide MS^2^PIP models, DeepLC, and Percolator into MS^2^Rescore thus allows substantial improved identification of novel epitopes from existing immunopeptidomics workflows.

## Introduction

The immune system is a complex, yet remarkable system that protects us from both invaders from outside the body, i.e. pathogens, as well as from inside the body, i.e. malignancies (1). Increased understanding of the immune system allowed for great medical achievements such as vaccination, which is currently available for over 29 diseases, enabled the eradication of smallpox, and prevents over 3 million deaths each year (2). However, many diseases such as *Mycobacterium tuberculosis* or malignancies lack effective vaccines due to improper T-cell activation. A key issue in developing effective vaccines for these diseases is the lack of accurately identified major histocompatibility complex (MHC)-presented epitopes or immunopeptides. These epitopes are presented on the cell surface and enable T-cells to discern healthy cells from infected or malignant cells. While much effort has recently been invested in accurate prediction of these epitopes *in silico* (3), these are mostly limited to viruses as these contain fewer potential protein antigens (4). Moreover, these tools are not yet sufficiently precise to confidently identify epitopes (5, 6). Therefore, experimental immunopeptidomics workflows, such as epitope detection through liquid chromatography-mass spectrometry (LC-MS), are still the best way to accurately identify these immunopeptides (7).

While immunopeptidomics workflows have been readily developed and applied (8), acquisition of immunopeptides through LC-MS suffers from some major problems. First, the acquisition of immunopeptide spectra is hampered due to the low abundance of immunopeptides and even more so of neo-epitopes. Infrequently occurring epitopes are still very challenging to identify through LC-MS, despite enrichment efforts and sample preprocessing (9). Second, in contrast to standard proteomics experiments where proteins are usually digested with trypsin before LC-MS, immunopeptides are captured through immune purification with antibodies followed by acidic elution resulting in mostly non-tryptic peptides. The non-tryptic nature of immunopeptides results in one less positive charge due to the missing arginine or lysine at the peptide’s C-terminus, causing many immunopeptides to be singly charged during MS acquisition. These singly charged peptides are much harder to analyze because, during fragmentation of the peptide, the charge resides on one of the fragments, leaving the other uncharged and thus lost (10). Moreover, most contaminants are singly charged as well, making identifications of immunopeptides much harder (11). The non-tryptic nature of immunopeptides hampers not only the acquisition but also the identification of immunopeptide spectra. To match each acquired spectrum with the peptide from which the spectrum originated, proteomics database search engines such as SEQUEST (12), X!Tandem (13), Andromeda (14), or PEAKS DB(15) generate *in silico* spectra for all potentially matching peptides. The complete list of peptides that could be in the sample is called the search space. It is important to note that spectra from peptides that are not included in this search space cannot be identified, even though they were acquired. In standard shotgun proteomics experiments, *in silico* tryptic digestion of relevant proteins yields a broad yet representable search space. In immunopeptidomics, however, the search space tends to be two orders of magnitude larger due to (i) seemingly random cleavage from the protein of origin, (ii) the variable length of MHC class I bound peptides, 8-11 amino acids, and MHC-II peptides, 6-24 amino acids (16) and (iii) the potential occurrence of conformational cis- and trans-spliced immunopeptides, which are non-linear peptides that originate from the same or different proteins, respectively (17). Additionally, sequence variants and non-canonical protein sequences are often considered as well, even further increasing the search space. Such a search space expansion leads to considerably more ambiguity between candidate peptide-spectrum matches (PSMs) (18), lower PSM scores, drastically elevated false discovery rate (FDR) score thresholds, and ultimately in fewer identified immunopeptides (19). Furthermore, because tryptic peptides have been the longtime standard in proteomics, search engines as well as bioinformatics tools that aid in identifying LC-MS spectra, are tailored towards tryptic peptides, making them less accurate or not applicable at all for immunopeptidomics.

The high need for neo- and xeno-epitope discovery led to the development of many bioinformatics tools to improve or validate identifications in immunopeptidomics. On the one hand, motif deconvolution tools have been developed that leverage binding motifs of immunopeptides to validate immunopeptide identifications. On the other hand, full pipelines have been developed to improve immunopeptide identification. For example, MHCquant (20), which is a recent computational workflow designed specifically for neo-epitope identification, and PEAKS DB (15). Even though PEAKS DB is not specifically designed for immunopeptides, it is highly interesting due to its *de novo* assisted database searches, which tend to work well for large search spaces. Even though these tools can help with immunopeptide identification, they do not use all available information, such as retention time and fragment ion intensity patterns. Previously, it has been proven that integrating retention time predictions in standard proteomics workflows can improve identification rates (21). Similarly, adding peak intensity predictions to post-processing tools such as Percolator can also improve identification rates drastically (22), which has already been proven to work for immunopeptides as well by efforts such as Prosit (23, 24). Similarly, tools such as DeepLC (25) and MS^2^PIP (26–28) can provide accurate retention time predictions and peak intensity predictions, respectively, to aid in post-processing. Indeed, when combined with Percolator, identification rates at a fixed FDR have been proven to substantially increase (22). However, currently, DeepLC and MS^2^PIP are solely trained on tryptic peptides. This absence of lysine and arginine at the C-terminus is less of a problem for DeepLC as the effect on retention time is small and is accounted for through feature encoding (29). However, this is not the case for MS^2^PIP, as alterations in peptide composition, as well as fragmentation patterns and labeling methods heavily alter peak intensity patterns (26). Therefore, we here present greatly improved MS^2^PIP models that include immunopeptides, and non-tryptic peptides in general. Moreover, we have integrated MS^2^PIP and DeepLC with Percolator into the free and open-source MS^2^Rescore software tool, which enables improved rescoring of peptide identifications from various proteomics search engines. Altogether, we show that well-adapted fragmentation spectrum and retention time predictions integrated into MS^2^Rescore drastically increase immunopeptide identification rates and outperform existing post-processing methods.

### Experimental Procedures

#### Training and evaluation of new MS^2^PIP spectrum prediction models for immunopeptides

To train and test new MS^2^PIP models, five publicly available immunopeptide data sets, and one publicly available chymotrypsin-digestion data set were downloaded from PRIDE Archive (30, 31). Similarly, for evaluation on representative unseen data, four distinct data sets were downloaded: (i) a data set containing HLA-I immunopeptides, (ii) a data set containing HLA-II immunopeptides, (iii) the data set with tryptic peptides that was previously used to evaluate the existing MS^2^PIP HCD models, and (iv) a data set containing chymotrypsin-digested peptide data. The corresponding ProteomeXchange project identifiers as well as the number of unique peptides and HLA patterns for each data set are listed in Supplementary Table S1.

As tandem mass spectrometry (MS2) fragmentation patterns are highly dependent on the instrument, instrument settings, fragmentation method, and any applied labeling methods (26), all MS^2^PIP train, test, and evaluation data must originate from experiments with the same experimental parameters. Unlabeled higher-energy collision-induced dissociation (HCD) data from Quadrupole-Orbitrap instruments was used, as this makes the newly trained models widely applicable and plenty of training data is readily available on public repositories. For each PRIDE Archive project, the raw mass spectrometry files were converted to Mascot Generic Format (MGF) files using ThermoRawFileParser (v1.3.4) (32). The corresponding identification files were converted to MS^2^PIP input files using custom Python scripts and were further filtered to retain unique combinations of peptide sequence, modifications, and precursor charge at 1% FDR. Next, all spectra were combined into one MGF file. The universal spectrum identifier (USI) (33) was used as unique identifier for each PSM to ensure reproducibility and a one-on-one mapping between peptide identifications and spectra. Data from each PRIDE Archive project was either used as train/test data or as evaluation data to ensure fully independent data sets, except for the chymotrypsin data, where the same project was used to provide both training/testing data (70%) and evaluation data (30%). This split was made after selecting unique peptide-modification-charge combinations, to ensure no overlap in samples between both splits.

Similarly to the 2019 MS^2^PIP models, new models were trained with the XGBoost machine learning algorithm (34). The Hyperopt (35), package (v0.2.5) was used in combination with four-fold cross-validation for hyperparameter optimization, allowing 400 boosting rounds and early stopping fixed at 10 rounds. Hyperparameters were optimized for each training data set separately, as well as for b- and y-ion models. All selected hyperparameters are shown in Supplementary Table S2. To evaluate each model, the Pearson correlation coefficient (PCC) was calculated between observed and predicted b- and y-ion peak intensities for each spectrum. The model performances were further analyzed by peptide length and precursor charge. Ultimately, three models were trained: the *immunopeptide* model solely trained on immunopeptides, the *immunopeptide-chymotrypsin* model trained on immunopeptides supplemented with chymotrypsin-digested peptides, and the *non-tryptic immunopeptide model* solely trained on non-tryptic immunopeptides. Ultimately, two models were integrated into MS^2^PIP: (i) the *immunopeptide* model and (ii) the *immuno-chymotrypsin* model. The former can be used for immunopeptide peak intensity predictions and the latter for tryptic and more general non-tryptic peptide predictions. For further analysis into rescoring immunopeptide PSMs, only the *immunopeptide* model was used.

To compare MS^2^PIP predictions with Prosit predictions, the same evaluation data sets were used as mentioned above. Prosit (v1.1.2) was downloaded from GitHub (https://github.com/kusterlab/prosit) and the hcd_hla and irt_prediction models were downloaded from Figshare (https://figshare.com/projects/prosit/35582). MS^2^PIP predictions were acquired for the general proteomics and chymotrypsin evaluation data with the immuno-chymotrypsin model, and for the HLA-I and HLA-II evaluation data with the immunopeptide model. Peptides that were not included in the Prosit output were filtered out of the MS^2^PIP predictions. The performance was measured in both PCC and spectral angle to ensure a thorough comparison. Only correlations for singly charged fragment ions were taken into account, as the newly trained MS^2^PIP models only predict intensities for these ions.

#### Evaluation of MS^2^Rescore on HLA class I peptides and comparison with Prosit rescoring

To validate the capacity of the new MS^2^PIP models to improve immunopeptide identification rates, the new models were implemented with DeepLC (v0.1.36) and Percolator (v3.5) into MS^2^Rescore. MS^2^Rescore calculates various meaningful features based on (i) the search engine output, (ii) the DeepLC-predicted and the observed retention times, and (iii) the MS^2^PIP-predicted and the observed MS2 peak intensities. These features are then passed to Percolator for PSM rescoring. Search engine features were selected based on the previous publication by Granholm et al. (36) and replicated for use with MaxQuant search results (37). MS^2^PIP features were used as first described by Silva et al (22). All features generated by MS^2^Rescore are listed in Supplementary Table S3.

MS^2^Rescore was validated on a large-scale HLA class I data set (38) which was also used to validate the recently published Prosit rescoring effort for immunopeptides (PXD021398) (24). This allows both an evaluation of the improved identification rates due to the new MS^2^PIP models, and a straight-forward comparison with Prosit rescoring. First, the msms.txt identification files for the projects’ two MaxQuant searches (alkylated and non-alkylated samples), the Prosit-rescored Percolator output files, and the raw mass spectrometry files were downloaded from PRIDE Archive. The mass spectrometry files were then further processed with ThermoRawFileParser (v1.3.4) (32) and the PSMs for each of the two MaxQuant searches were rescored separately. Two rescoring methods were evaluated: (i) Using only search engine features, replicating a normal Percolator run, and (ii) using the full MS^2^Rescore feature set, including search engine-, MS^2^PIP-, and DeepLC-features. Additionally, these rescoring methods were compared with the original MaxQuant results and with the downloaded Prosit-rescoring results.

Each rescoring method was evaluated at varying FDR thresholds in terms of identification rate and number of unique identified peptides. The contribution of the different feature sets in MS^2^Rescore was visualized using Percolator’s model weights, and the distributions of retention time difference and MS^2^PIP prediction correlations were compared between decoy PSMs, accepted target PSMs, and rejected target PSMs.

Additionally, as reported by Wilhelm et al. (24), sequence motif patterns for HLA pattern C*12:03 were further analyzed with GibbsCluster (v2.0) (39), for the gained and lost peptides compared to rescoring with only search engine features.

#### Evaluation of MS^2^Rescore across collision energy settings and peptide abundances

To further analyze MS^2^Rescore performance for various experimental collision energy settings, replicate LC-MS/MS runs were performed on HL60 cells at collision energy values of 25, 27, 30, 32 and 35 NCE (Supplementary Methods). The resulting spectra were searched with the Andromeda search engine (MaxQuant v1.6.14.0) against the human UniProtKB-SwissProt (14-09-2020; 20 388 sequences, Taxonomy ID 9606) database without any enzyme specificity. A minimal peptide length of seven amino acids was required. Oxidation (M) was set as variable modification with a maximum of three modifications per peptide. Mass tolerances were set at 5 ppm and 20 ppm for MS1 and MS2 spectra respectively. False discovery rate was kept at 100% with the use of a decoy strategy for downstream rescoring with (i) only search engine features, and (ii) the full MS^2^Rescore feature set. Furthermore, for all LC-MS/MS runs at all collision energy settings, precursor intensities were obtained from the MaxQuant msms.txt file to assess any differences in the performance of MS^2^Rescore between low and high abundant peptides.

#### Evaluation of MS^2^Rescore on HLA class II peptides

To validate MS^2^Rescore for HLA class II peptides, another set of raw mass spectrometry files were downloaded from PRIDE Archive (PXD015408). As the uploaded search engine results were already filtered at 5% FDR, the spectra were re-analyzed with PEAKS DB (v10.5) (15) with the same search parameters that were used in the original publication, i.e. no enzyme specificity, precursor error tolerance of 10 ppm, fragment ion tolerance of 0.01 Da, with oxidation (M), deamidation (NQ) and trioxidation (C) as variable modifications, searched against the UniProtKB-SwissProt database (01-2021, 22 235 sequences, Taxonomy ID 10090). The mzIdentML identification file as well as the corresponding MGF files were exported from PEAKS DB and were rescored with MS^2^Rescore with (i) only search engine features, and (ii) the full MS^2^Rescore feature set, as described above for the evaluation on HLA class I peptides.

#### General data processing and data visualization

All plots, unless specified, were generated in Jupyter notebooks (v6.4.0) using Python (v3.8.3) with the Matplotlib (v3.4.2) (40), Seaborn (v0.11.0) (41), UpSetPlot (v0.6.0), and spectrum_utils (v0.3.5) (42) libraries.

## Results

### Newly trained MS^2^PIP models accurately predict immunopeptide spectrum peak intensities

In order to improve the identification rate of immunopeptides by leveraging peak intensity predictions, new models for MS^2^PIP were trained specifically for immunopeptides. Despite using different training set compositions, all newly trained models drastically improve predictions for both HLA-I and HLA-II data in comparison with the tryptic 2019 HCD model (Figure 1A). Surprisingly, even for standard tryptic shotgun proteomics data the predictions from the new models are slightly better, largely due to the portion of tryptic peptides within the immunopeptide training data. Indeed, when these peptides are left out of the training data, accuracy drops in comparison with the 2019 HCD model. While both immunopeptide models are well suited to predict peak intensities for tryptic- and immunopeptides, the performance on chymotrypsin-digested peptides is not as high. Only when chymotrypsin-digested peptides are included in the training data, performance on chymotrypsin data increases. This difference in performance between models trained with or without chymotrypsin-digested peptides, is mainly seen for longer peptides and peptides with higher precursor charge states (Supplementary Figure S1). As models that are solely trained on immunopeptides have not seen longer peptides with higher charge-states, these models can not accurately model how these will behave in the mass spectrometer. Thus, even though immuno- and chymotrypsin-digested peptides are both considered non-tryptic, they are still very different. Overall, immunopeptide peak intensity predictions are drastically improved by all the newly trained models, with the immunopeptide model showing the highest accuracy (median PCC of 0.94). The exact median PCCs are listed in Supplementary Table S4. Examples of a prediction with median PCC values with the immunopeptide model and the corresponding, less accurate, 2019 model prediction are shown in Figure 1B and Figure 1C.

**Figure 1.**
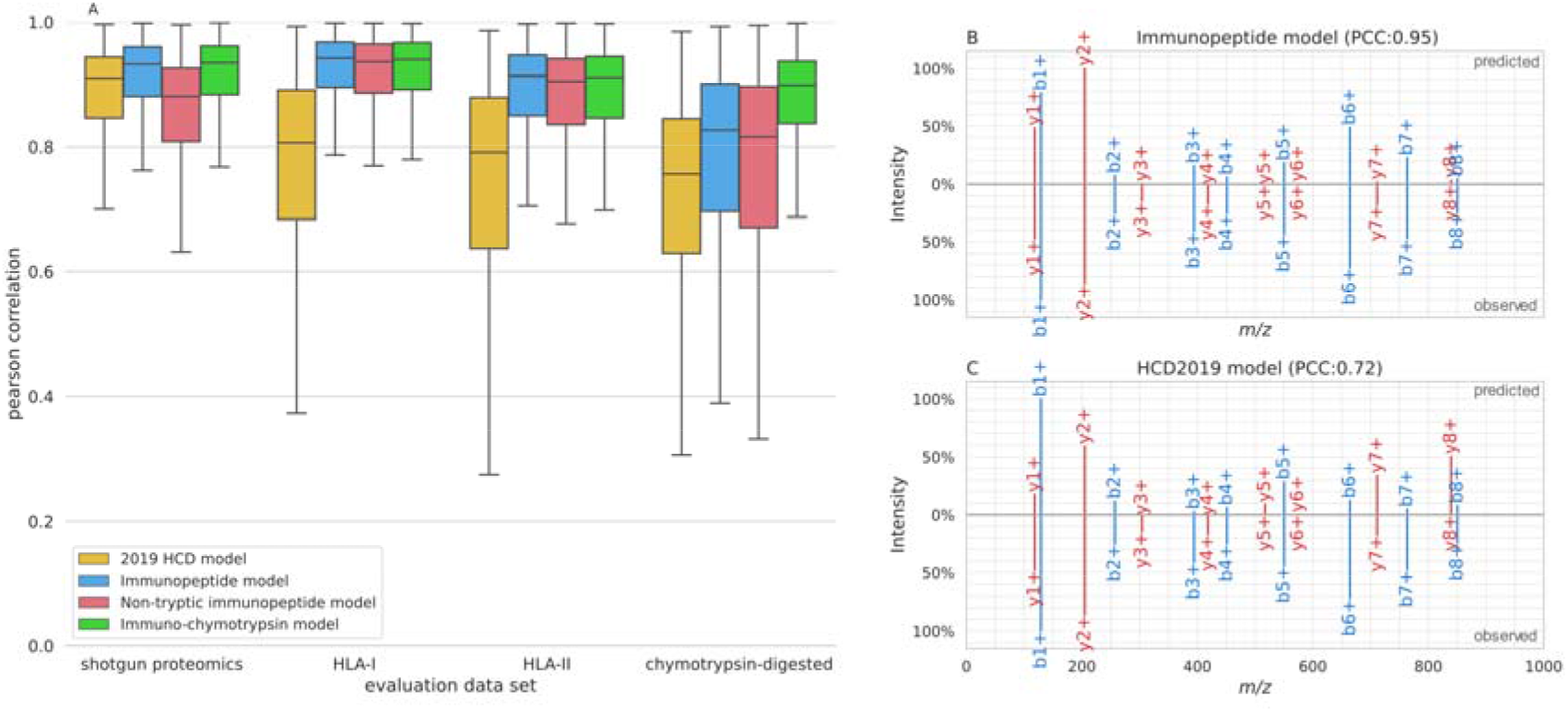
Performance evaluation of the new non-tryptic MS^2^PIP models. Boxplots comparing Pearson correlation distributions between predicted and observed peak intensity values on spectrum level for the 2019 HCD model, immunopeptide model, non-tryptic immunopeptide model, and immuno-chymotrypsin model (A). Performance is evaluated on general tryptic proteomics data, HLA-I immunopeptide data, HLA-II immunopeptide data, and chymotrypsin data. Predictions for immunopeptide “KQHGVNVSV” by new immunopeptide MS^2^PIP model (top) compared to observed MS2 spectrum (mzspec:PXD005231:20160513_TIL1_R1:scan:10909) (bottom) (B). This peptide was selected for visualization as its PCC lies close to the median for the immunopeptide model on all HLA-I evaluation data. Predictions by the 2019 HCD model for the same immunopeptide (C) as visualized in B.

### MS^2^Rescore drastically improves immunopeptide identification rates

Ultimately, the goal of these newly trained models is to improve immunopeptide identification rates by providing more accurate peak intensity predictions. Therefore, (i) identification results without rescoring, (ii) rescoring with solely search engine features (replicating a normal Percolator run), (iii) rescoring with MS^2^Rescore (including DeepLC and the new MS^2^PIP models), and (iv) rescoring with the recently published Prosit models were compared in terms of the total amount of identifications as well as the number of unique identifications based on sequence. Overall, rescoring with both MS^2^Rescore and Prosit substantially improved the spectrum identification rate in comparison with rescoring with search engine features alone or not rescoring, and this at both 1% and 0.1% FDR. Indeed, MS^2^Rescore achieves an identification rate of 11,1%, out of 18 million spectra, compared to 7.6% for traditional rescoring (an increase of 46%), and only 1.9% for the MaxQuant search results, all at 1% FDR (Figure 2A-B). Moreover, 83% of the identified spectra at 1% FDR are retained when restricting the threshold to 0.1% FDR. Thus, providing peak intensities and retention time predictions to Percolator substantially increases the number of identified immunopeptides. This is clearly illustrated by analyzing the Percolator weights for each separate feature as well as the combined absolute weights for search engine features, MS^2^PIP features and DeepLC features (Supplementary Figure S2). Similarly, the number of unique identified immunopeptides increases by 36% when adding MS^2^PIP and DeepLC Features for the 1% FDR, and even more so for 0.1% FDR where the number of unique identified peptides reaches nearly 300% of the number of traditional Percolator identification results (Figure 2C-F). These gains are consistent across all 95 HLA class I alleles included in the data (Supplementary Figure S3-4), showing that the newly trained MS^2^PIP model, and therefore MS^2^Rescore, is generalizable across different HLA types. In fact, MS^2^Rescore allows for a substantial increase in identification rate for HLA types with initially fewer identifications (e.g., A0101, A0204, B4402…), indicating that MS^2^Rescore especially improves the peptide identification coverage for harder-to-identify HLA alleles.

**Figure 2.**
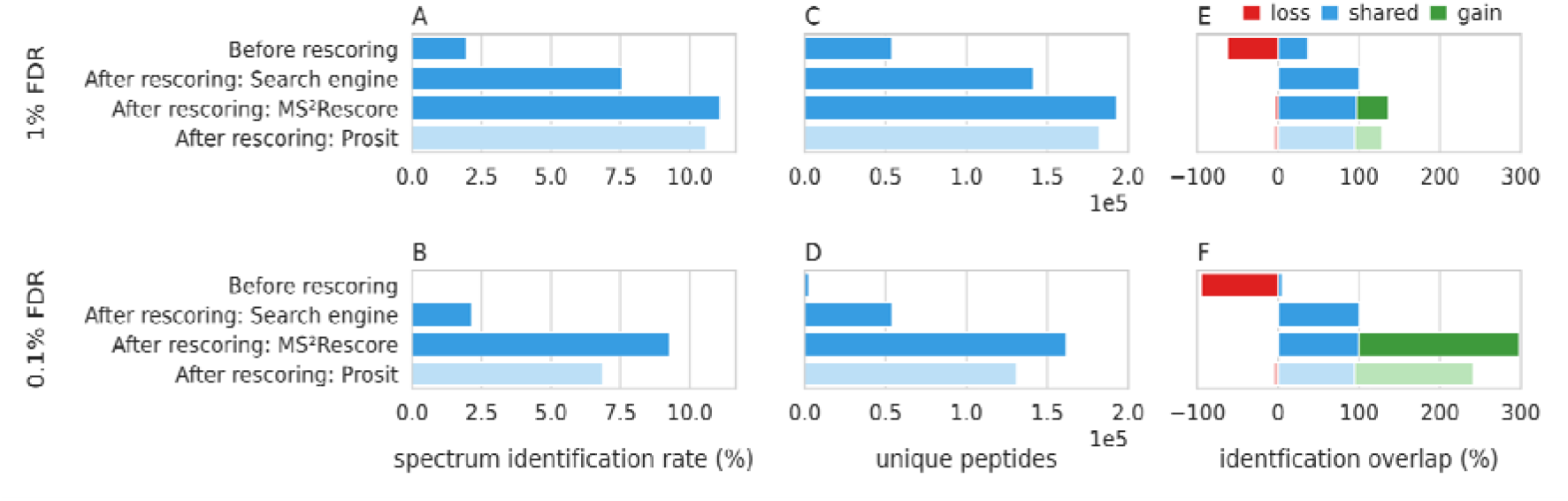
Percentage of identified spectra and unique identified peptides using different rescoring methods (PXD021398). Bar charts showing the spectrum identification rate out of 18.375.659 spectra (A, B), showing the total number of unique identified peptides in terms of sequence (C, D), and showing the shared (blue), gained (green) and lost (red) number of unique (by sequence) identified immunopeptides in relation to rescoring with only search engine features. for the 1% FDR (E) and 0.1%FDR (F) threshold. All results are shown for the 1% FDR (A, C, E) and 0.1% FDR (B, D, F) thresholds.

The power of providing these predictions to Percolator is further illustrated when visualizing the distributions for decoy PSMs, rejected target PSMs, and accepted target PSMs. Indeed, the distributions for decoy and rejected target PSMs are highly similar for both the retention time error as well as the PCC, while the accepted target PSMs accumulate around low retention time errors and high PCCs (Figure 3A-B). The accepted target PSMs are clearly separable from the decoy and rejected target PSMs using only the PCC and retention time error distributions (Figure 3C-D). Furthermore, while both metrics correlate with the search engine score, a large amount of decoy and rejected target PSMs can only be separated from the target PSMs by also including PCC or retention time error information (Figure 3E-F). This clearly illustrates how Percolator achieves its much-improved separation between true and false target PSMs when provided with peak intensity and retention time prediction features. Furthermore, PSMs that previously would have been incorrectly accepted below a 1% FDR because of a high search engine score alone, are now rejected due to a low PCC, a high retention time error, or both. This most likely accounts for the small percentage of identified peptides that are lost after rescoring.

**Figure 3.**
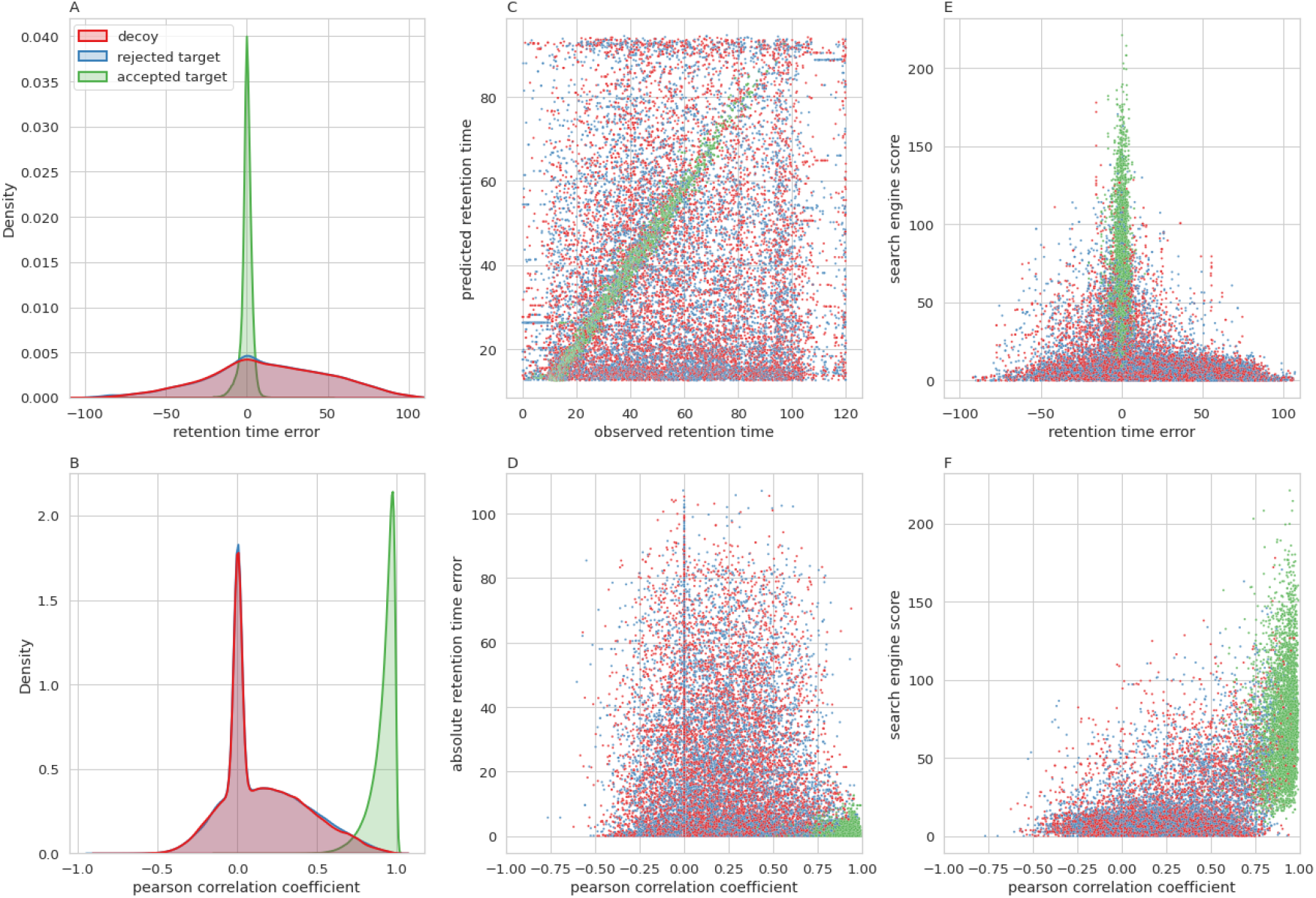
Distributions of MS^2^PIP-, DeepLC and search engine-based features. Density plots showing the distribution of the smallest retention time error between observed and predicted retention time (A) and showing the Pearson correlation between observed and predicted peak intensities (B) for each unique combination of sequence and modification for decoys (red) rejected targets, q value ć 0.01(blue) and accepted targets, q value ć 0.01 (green) (note the rejected targets distribution coincides with the decoy distribution). Scatterplots showing the relation between observed and predicted retention time (C), between Pearson correlation and retention time error (D), between retention time error and search engine score (E), and between Pearson correlation and search engine score (F) for decoys (red), rejected targets (blue), and accepted targets (green) for all PSMs from “GN20170531_SK_HLA_C0102_R1_01”. Note that all retention time errors are the smallest retention time error for each precursor (see rt_diff_best table S3), all Pearson correlation coefficients are calculated for each full log2 transformed spectrum, and all zero values are caused by the fact that either the observed or predicted intensities for a given ion type are all zero.

### MS^2^Rescore outperforms the current state-of-the-art

The integration of MS^2^PIP-, DeepLC-, and search engine-based features in MS^2^Rescore has proven to substantially increase the identification rate of immunopeptides, and furthermore, outperforms the recently published Prosit rescoring method (24). In comparison with Prosit, MS^2^Rescore gains 5% and 35% more identifications at 1% FDR and 0.1% FDR, respectively. This trend continues for the number of unique identified peptides with a respective increase of 8% and 57% (Figure 2). Indeed, over 32 000 unique peptides were identified below the 0.1% FDR threshold by MS^2^Rescore while Prosit rescoring only identified these peptides at the 1% FDR threshold, highlighting the gain in confidence in terms of unique identified immunopeptides (Supplementary Figure S5). MS^2^Rescore thus substantially increases the identification rate, especially for more stringent FDR thresholds. Peptides for which Prosit cannot predict MS2 peak intensities, i.e. unmodified (C), cysteinylated (C), and acetylation (N-terminus), were left out of the Prosit rescoring output. That is why MS^2^Rescore includes more PSMs in the unfiltered data set at 100% FDR, especially for the noIAA sample (Supplementary Figure S6A-B). For a second, thorough comparison, these PSMs were left out of the MS^2^Rescore output as well. However, this seemed to have a rather negligible impact on the number of identifications at 1% and 0.1% FDR (Supplementary Figure S6C-D), and thus the difference in identification rates cannot be attributed to these filtered peptides.

Furthermore, a comparison of the peptide spectrum prediction accuracy of the newly trained MS^2^PIP models and Prosit indicates that higher performance of MS^2^Rescore cannot be attributed to improved peak intensity predictions. Indeed, depending on the correlation metric that is used, either MS^2^PIP or Prosit performs slightly better than the other on the evaluation data (Supplementary Figure S7). It is important to note, however, that for Prosit the correct CE value should be selected for optimal performance. To determine if the difference in rescoring between MS^2^Rescore and Prosit is driven by a difference in features, all MS^2^Rescore features that do not have a close counterpart in Prosit rescoring were removed for a separate run. This includes all DeepLC retention time features and various search engine features (Supplementary Table S3). As MS^2^Rescore and Prosit both include various similar peak intensity prediction features, these were retained. With this reduced feature set, the performance of MS^2^Rescore is slightly lower than for Prosit rescoring (Supplementary Figure S6A-D), which confirms that the retention time features and additional search engine features result in improved MS^2^Rescore performance over Prosit rescoring.

Similarly to Prosit, the identified sequence motif for HLA type C*12:03 was highly similar to the motif reported in the original publication of the data set (24, 38), while the peptides that were removed by MS^2^Rescore compared to search engine rescoring showed quite different, less conserved sequence motifs (Supplementary Figure S8).

### MS^2^Rescore generalizes well across collision energy settings and peptide abundances

Because MS^2^PIP does not account for CE in its predictions, MS^2^PIP and consequently MS^2^Rescore could potentially be biased towards spectra obtained with certain CE values. Therefore, search results of replicate mass spectrometry runs with CE settings varying from 25 to 35 were post-processed with MS^2^Rescore. For each CE value, MS^2^Rescore shows a significant increase in identification rate. However, for larger (less optimal) CE values, the overall identification rate decreases (Supplementary Figure S9A). This is most likely due to a reduced quality in fragmentation spectra, which is reflected in the decreased explained ion current, and in line with the b- and y-ion MS^2^PIP PCC distributions for accepted PSMs when using suboptimal CE values (Supplementary Figure S9D-F). Most interestingly, the relative gain in unique identified peptides for MS^2^Rescore increases for higher (and therefore less optimal) CE values, approaching a 60% increase for CE 35, by slightly shifting the feature weights away from fragmentation features in favor of DeepLC retention time features (Supplementary Figure S9B-C). Consequently, MS^2^Rescore is able to recover peptides that would otherwise be lost due to lower-quality fragmentation spectra.

A similar effect is observed for low abundant peptides. Indeed, the largest relative gain achieved by MS^2^Rescore in terms of number of unique identified peptides is seen for the lowest precursor intensities (Supplementary Figure S10A-B), where traditional rescoring fails to recover most identifications. MS^2^Rescore is thus not only able to increase the amount of identifications for immunopeptides in general, it can recover peptides previously lost due to low precursor intensities and thus lower-quality spectra or nonoptimal instrument settings.

### MS^2^Rescore is unbiased to different HLA classes

To evaluate the performance of MS^2^Rescore on HLA class II peptides, another publicly available data set was re-analyzed. However, while for the HLA class I data set human immunopeptides and MaxQuant search engine (14) results were used, for this HLA class II data set, mouse data was searched with PEAKS DB (15). As was the case for HLA class I peptides, MS^2^Rescore significantly increases the identification rate for HLA class II peptides with 15% and 57% for the 1% and 0.1% FDR threshold, respectively (Supplementary Figure S11). These increases are, however, slightly lower than for the HLA class I data set. Moreover, where previously conventional rescoring showed a significant increase in comparison to search engine rescoring, here the gain in comparison with no rescoring is lower for both identification rate as well as number of unique identified peptides. This is likely due to (i) the less extensive search engine features that are calculated for the PEAKS DB pipeline in MS^2^Rescore (Supplementary Table S3), and (ii) to the fact PEAKS DB is likely better equipped to identify immunopeptides than MaxQuant due to its *de novo*-assisted database search (15). Nevertheless, the full MS^2^Rescore feature set, including peak intensity and retention time predictions, still results in a significantly higher identification rate. Altogether, these results show that MS^2^Rescore generalizes well across HLA class I and class II immunopeptides, across different species, and can boost the performance from different search engines.

## Discussion

By training new peak intensity prediction models, we were able to greatly enhance immunopeptide identification rate through PSM rescoring. While all newly trained models greatly enhance peak intensity predictions for immunopeptides, the model trained solely on immunopeptides performed best. Even though the immuno-chymotrypsin model contained the same immunopeptide train set, the addition of the chymotrypsin-digested peptides did lower the performance slightly. Similarly, not including chymotrypsin-digested peptides in the training data resulted in lower accuracies for the chymotrypsin-digested peptides. Indeed, immunopeptides are generally much smaller and consequently carry a lower charge state. Therefore, immunopeptide trained models are not able to predict the behavior of longer and higher charged peptides in the mass spectrometer. While both immuno- and chymotrypsin-digested peptides are considered non-tryptic, their properties can be very different, leading to reduced accuracy of peak intensity prediction models when applied on a different type of non-tryptic peptides. Therefore, a general non-tryptic peak intensity prediction model seems not ideal, as it could lower performance for subsets like immunopeptides. Surprisingly, while immuno- and chymotrypsin-digested peptides are antagonistic, immuno- and tryptic peptides seem synergistic in terms of training data. This comes as no surprise as almost 50% of the immunopeptide training data consists of tryptic peptides. Immunopeptides are thus not necessarily non-tryptic. However, the actual occurrence of tryptic peptides in immunopeptidomics samples is most likely much lower. This unrepresentatively sized tryptic portion most likely originated from the tryptic bias in current immunopeptidomics workflows. Indeed, in previous studies, tryptic MHC peptide coverage could rise to 70% (43). By training new, non-tryptic models of MS^2^PIP, we take a first step in decreasing this tryptic bias, to ultimately be able to analyze an unbiased immunopeptide landscape.

Moreover, by integrating the new immunopeptide model with retention time predictions and search engine features into MS^2^Rescore, we greatly enhanced the ability of Percolator to rescore immunopeptide PSMs, resulting in a much-improved immunopeptide identification workflow. Furthermore, rescoring drastically increases the number of unique identified peptides, which is of crucial importance for the discovery of potential neo-epitopes for cancer vaccination or xeno-epitopes for anti-bacterial, and to a lesser extent, anti-viral vaccines. Moreover, while previously almost no identifications were found at a more confident 0.1% FDR threshold, MS^2^Rescore allows a lowering of the FDR threshold to 0.1%, while retaining 83% of the peptides identified at 1% FDR. This illustrates the large increase in confidence of the identified PSMs MS^2^Rescore introduces. Besides the increase in both PSM confidence and identification rate, MS^2^Rescore has shown to be unbiased with regard to HLA patterns and CE settings. Most importantly, the relative identification gain introduced by MS^2^Rescore is even larger for HLA patterns that initially had fewer identifications, showing that MS^2^Rescore is able to increase the view on the immunopeptide landscape for traditionally harder-to-identify HLA patterns. Moreover, MS^2^Rescore is able to recover peptide identifications that would have been lost due to lower quality spectra by making use of DeepLC retention time features, and can therefore recover substantial additional identifications for low abundant peptides. This potentially enables the recovery of biologically relevant neo- or xeno-epitopes that occur less frequently in the sample. Furthermore, MS^2^Rescore is able to gain immunopeptide identifications regardless of the search engine used, for both HLA class I and class II peptides, and across different species.

Additionally, MS^2^Rescore with DeepLC and the new immunopeptide MS^2^PIP models shows an improved identification rate over the recently published Prosit effort, especially for lower FDR thresholds. As Prosit has shown to provide more accurate predictions compared to previous MS^2^PIP models, it is unlikely that MS^2^Rescore’s higher performance can be attributed to superior peak intensity predictions. Indeed, the peak intensity prediction accuracies of the new MS^2^PIP models and of Prosit are highly similar for immunopeptides (Supplementary Figure S7) even when Prosit has been optimized for the right CE. These negligible differences in peak intensity prediction correlations are therefore likely not the reason for the higher performance of MS^2^Rescore in favor of Prosit. Instead, it is more likely that the main difference in rescoring performance is the result of the generation of more relevant MS^2^PIP-, DeepLC and search engine-derived features. Indeed, when the majority of the search engine features and all DeepLC retention time features were omitted, reflecting the more limited Prosit feature set, the performance of MS^2^Rescore drops as well (Supplementary Figure S6). By providing a more extensive feature set, MS^2^Rescore creates a unique feature space that allows Percolator to separate true from false identifications much better than when provided with limited features without retention time or peak intensity information (Figure 3). The combination of all these calculated features is therefore likely to be driver of MS^2^Rescore’s superior performance.

MS^2^Rescore is freely available under the permissive Apache 2.0 open-source license on GitHub (https://github.com/compomics/ms2rescore) and can easily be installed locally through the cross-platform PyPI Python package, as well as with a standalone windows install script. Both a command line interface and a graphical user interface are available, various identification files from different search engines are accepted, and both MS^2^PIP and DeepLC can handle a variety of modifications, eliminating the need to filter identification files before rescoring. Altogether, these new models show great promise to greatly extend the immunopeptide landscape in existing and future immunopeptidomics experiments.

## Abbreviations

CE: collision energy
Da: dalton
FDR: false discovery rate
HCD: higher-energy collision-induced dissociation
HLA: human leukocyte antigen
LC-MS: liquid chromatography-mass spectrometry
MS2: tandem mass spectrometry
MHC: major histocompatibility complex
MGF: mascot generic format
PCC: Pearson correlation coefficient
PSM: peptide-spectrum match
USI: universal spectrum identifier

## Acknowledgments

R.B. acknowledges funding from the Vlaams Agentschap Innoveren en Ondernemen under project number HBC.2020.2205.; R.G. received funding from the Research Foundation Flanders (FWO) [1S50918N]. S.D. and L.M. acknowledge funding from the European Union’s Horizon 2020 Programme (H2020-INFRAIA-2018-1) [823839]; L.M. acknowledges funding from the Research Foundation Flanders (FWO) [G028821N] and from Ghent University Concerted Research Action [BOF21/GOA/033]. A.D. received funding from the Research Foundation Flanders (FWO) [1SE3722]. NanoLC-MS/MS instruments were supported by the French Proteomic Infrastructure (ProFI FR2048; ANR-10-INBS-08-03).

## Author contributions

AD, SD, LM, and RG conceived the project. AD developed the new MS^2^PIP models, carried out the implementation in MS^2^Rescore, and performed all validations. RG developed the initial MS^2^Rescore Python package. RB assisted with the integration and validation of the DeepLC models into MS^2^Rescore. AH and CC conducted the collision energy ramp experiments. AD and RG analyzed the results. AD and RG wrote the manuscript, and all authors contributed to the final version.

## Conflict of interest

The authors declare that they have no conflict of interest.

## Data availability

MS^2^Rescore is available at https://github.com/compomics/ms2rescore. All additional code used in this work is available at compomics/ms2rescore-immunopeptidomics-manuscript (github.com). The data used for training and evaluation of the newly trained models, the models themselves as well as the MS^2^Rescore output is available on Zenodo at https://doi.org/10.5281/zenodo.6532013. Additional mass spectrometry proteomics data have been deposited to the ProteomeXchange Consortium via the PRIDE (31) partner repository with the dataset identifier PXD033868.

## Notes

https://doi.org/10.5281/zenodo.6532013

### Competing Interest Statement

The authors have declared no competing interest.

### Summary of Updates

The manuscript has been updated with a more in depth analysis of the HLA class I rescoring. Supplementary Figure S3 & S4 have been added to show the performance of MS^2^Rescore across different HLA types while the new Supplementary Figure 8 shows that the peptides gained for HLA type C*12:03 by rescoring nicely fit the established pattern for this HLA type from the original paper. To compare with current state-of-the-art, Prosit, we evaluated the overlap in unique identified peptides between MS^2^Rescore and Prosit (Supplementary Figure S5). Furthermore, we added a peak intensity comparison between Prosit and the newly trained MS^2^PIP models (Supplementary Figure S7) and we, temporarily, removed features from MS^2^Rescore that were not included in Prosit rescoring to establish which features drive the difference between both rescoring methods (Supplementary Figure S6). Next, we have evaluated the performance of MS^2^Rescore across different collision energy settings (Supplementary Figure S9), peptide abundances (Supplementary Figure S10), and we have added a results section on rescoring HLA class II mouse data that was originally searched with PEAKS DB (Supplementary Figure S11). Lastly, we have made some small figure changes to Figure 1, Figure 2 and Figure 3 as well as Supplementary Figure S2 and Dr. Christine Carapito and Aurélie Hirschler who have conducted the collision energy ramp experiments were added to the authors list of this manuscript.

