## Supplementary data for "MS^2^Rescore: Data-driven rescoring dramatically boosts immunopeptide identification rates"

### Supplementary methods

**Acquisition and identification of additional LC-MS/MS runs for the assessment of the generalizability of MS²Rescore across collision energy settings and peptide abundances**

In total 600 million HL60 cells were lysed with 20mM Tris-HCl, 150 mM NaCl, 0,25% sodium deoxycholate, 1mM EDTA pH8, 0,2mM iodoacetamide, 1mM PMSF, Roche Complete Protease Inhibitor Cocktail, 0,5 % NP 40, PBS, pH 7,4. The lysate was centrifuged at 21 000g for 30 minutes at 4°C. MHC-peptide-complexes were captured on CNBr-activated sepharose 4B beads (Cytivia) linked to an antibody. Following binding, beads were washed several times with 3 buffers (150 mM NaCl, 20 mM Tris-HCl, pH7.4; 400 mM NaCl, 20 mM Tris-HCl, pH 7.4; and 20 mM TrisHCl, pH 8.0) and bound complexes were eluted in 0.1M acetic acid. Eluted HLA peptides and the subunits of the HLA complexes were desalted using a C18 Macro Spin column (Harvard Apparatus) according to the manufacturer’s protocol. Finally, HLA peptides were purified from the MHC-I complex after the elution with 25% ACN, 0.1% TFA. Samples were evaporated under vacuum and resuspended in H2O with 0.1% FA.

NanoLC-MS/MS analyses were performed on a nanoAcquity UltraPerformance Liquid Chromatography device (Waters Corporation) coupled to a quadrupole-Orbitrap mass spectrometer (Q-Exactive HF-X, Thermo Fisher Scientific). Peptide separation was performed on an ACQUITY UPLC® Peptide BEH C18 Column (250 mm x 75 µm with 1.7 µm diameter particles) and an ACQUITY UPLC® M-Class Symmetry® C18 Trap Column (20 mm x 180 µm with 5 µm diameter particles; Waters). The solvent system consisted of 0.1% FA in water (solvent A) and 0.1% FA in ACN (solvent B). Samples (400 ng) were loaded into the enrichment column over 3 minutes at 5 μL/min with 99% of solvent A and 1% of solvent B. Chromatographic separation was conducted with the following gradient of solvent B: from 1 to 25% over 90 min, from 25 to 90% over 1 min. The MS capillary voltage was set to 2 kV at 250°C. The system was operated in a data-dependent acquisition mode with automatic switching between MS (resolution of 120 000 at 200 m/z, automatic gain control fixed at 3 x 10^6^ ions and a maximum injection time set at 80 milliseconds) and MS/MS modes (resolution of 30 000 at 200 m/z, automatic gain control fixed at 1 x 10^5^, and the maximal injection time set at 240 milliseconds) to acquire high-resolution MS/MS spectra. The ten most abundant peptides were selected on each MS spectrum for further isolation and higher energy collision dissociation, excluding unassigned, monocharged and superior to seven times charged ions. A ramp of five different collision energy settings were applied (25, 27, 30, 32 and 35 NCE ) and each setting was tested in triplicate. A solvent blank injection was performed after each sample to minimize carry-over.

### Supplementary tables

#### Supplementary Table S1

List of all PRIDE Archive projects used for training, testing, and evaluating new MS²PIP models. Unique peptides denote the total number of unique peptides in terms of sequence, modifications, and precursor charge used in the training set for each project with the number of unique HLA allele types, both class I and II for each project.

| PRIDE Archive project | Unique peptides | Peptide type | HLA-alleles(HLA-I/HLA-II) | Publication |
| --- | --- | --- | --- | --- |
| **Train / test** | | | | |
| PXD012308 | 61 590 | Immunopeptide (HLA-II) | -/64 | (Racle *et al*, 2019) |
| PXD006939 | 120 427 | Immunopeptide (HLA-I/II) | 37/28 | (Chong *et al*, 2018) |
| PXD009925 | 17 339 | Immunopeptide (HLA-I) | 84/- | (Gfeller *et al*, 2018) |
| PXD000394 | 134 891 | Immunopeptide (HLA-I) | 28/- | (Bassani-Sternberg *et al*, 2015) |
| PXD004894 | 129 011 | Immunopeptide (HLA-I/II) | 14</0<* | (Bassani-Sternberg *et al*, 2016) |
| PXD010154 (70%) | 60 598 | Chymotrypsin-digested |  | (Wang *et al*, 2019) |
| **Evaluation** | | | | |
| PXD005231 | 12 534 | Immunopeptide (HLA-I) | 33/- | (Bassani-Sternberg *et al*, 2017) |
| PXD020011 | 12 745 | Immunopeptide (HLA-II) | 16/- | (Marino *et al*, 2020) |
| PXD010154 (30%) | 25 570 | Chymotrypsin-digested |  | (Wang *et al*, 2019) |
| PXD008034 | 35 212 | General proteomics data |  | (Gravina *et al*, 2018) |

*Only 5 out of the 25 patients were HLA typed.

#### Supplementary Table S2

| Model | Eta | Max depth | Grow policy | Max leaves | Min child weight | Gamma | Lambda | Alpha | Colsample by tree | Sub-sample |
| --- | --- | --- | --- | --- | --- | --- | --- | --- | --- | --- |
| Immuno-chymotrypsin (b-ions) | 0.08060612330262913 | 18 | Lossguide | 117 | 500 | 0.031142279181653326 | 0.2724553826622634 | 3.4 | 0.891381182690278 | 0.7 |
| Immuno-chymotrypsin (y-ions) | 0.047107785048838 | 18 | Lossguide | 490 | 4 | 0.37528441949267444 | 0.35150807248415 | 3.3 | 0.6122042447952851 | 0.6 |
| Immunopeptide (b-ions) | 0.09263630381479264 | 17 | Lossguide | 131 | 16 | 0.6048882172751935 | 0.9332236183206803 | 4.6 | 0.9898165069470042 | 0.7 |
| Immunopeptide (y-ions) | 0.0594145790364741 | 17 | Lossguide | 302 | 3 | 0.03338151150211477 | 0.4430375595950531 | 4.5 | 0.9389820388602939 | 0.7 |
| Non-tryptic immunopeptide (b-ions) | 0.05754126608420542 | 16 | Lossguide | 356 | 85 | 0.35987834628709725 | 0.7049467859206121 | 3.5 | 0.7002269364273864 | 0.6 |
| Non-tryptic immunopeptide (y-ions) | 0.08443065926710414 | 18 | Lossguide | 6 | 7 | 0.38091488564428677 | 0.6137299313065955 | 4.3 | 0.7958195937743725 | 1.0 |

The optimal hyperparameters for each new b- and y-ion MS²PIP model, as determined during hyperparameter optimization.

#### Supplementary Table S3

Percolator rescoring features generated by MS²Rescore from MaxQuant search results.

| Feature name | Description |  |
| --- | --- | --- |
| **Search engine features (MaxQuant integration)** | | **No equivalent in Prosit rescoring** |
| ln_ms2_ion_current | Natural logarithm of the summed intensity of all annotated fragment ions | x |
| ln_cterm_ion_current_ratio | Natural logarithm of the summed intensity of the annotated y-ions, divided by that of all annotated fragment ions | x |
| ln_nterm_ion_current_ratio | Natural logarithm of the summed intensity of the annotated b-ions, divided by that of all annotated fragment ions | x |
| ln_explained_ion_current | Natural logarithm of the summed intensity of the annotated fragment ions, divided by that of all fragment ions | x |
| stdev_error_top7 | Standard deviation of the mass errors of the seven most intense annotated fragment ions | x |
| sq_mean_error_top7 | Squared mean of the mass errors of the seven most intense annotated fragment ions | x |
| mean_error_top7 | Mean of the mass errors of the seven most intense annotated fragment ions | x |
| Charge7 | One-hot encoded precursor charge state (1 if charge is 7, else 0) |  |
| Charge6 | One-hot encoded precursor charge state (1 if charge is 6, else 0) |  |
| Charge5 | One-hot encoded precursor charge state (1 if charge is 5, else 0) |  |
| Charge4 | One-hot encoded precursor charge state (1 if charge is 4, else 0) |  |
| Charge3 | One-hot encoded precursor charge state (1 if charge is 3, else 0) |  |
| Charge2 | One-hot encoded precursor charge state (1 if charge is 2, else 0) |  |
| Charge1 | One-hot encoded precursor charge state (1 if charge is 1, else 0) |  |
| absdM | Absolute difference between theoretical and observed precursor mass |  |
| enzInt | Number of missed enzymatic cleavages |  |
| dM | Difference between theoretical and observed precursor mass |  |
| PepLen | Peptide length |  |
| Mass | Precursor mass |  |
| ChargeN | Precursor charge state |  |
| RawModLocProb | Andromeda modification localization probability | x |
| RawDeltaScore | Difference in Andromeda score with next lower-ranking PSM | x |
| RawScore | Andromeda score | x |
| **Search engine features (PEAKS DB integration)** | |  |
| PEAKS:peptideScore | PEAKS scores |  |
| calculatedMassToCharge | Calculated mass |  |
| experimentalMassToCharge | Experimental mass |  |
| dM | Difference between theoretical and observed precursor mass |  |
| absdM | Absolute difference between theoretical and observed precursor mass |  |
| Peptide_length | Peptide length |  |
| Charge7 | One-hot encoded precursor charge state (1 if charge is 7, else 0) |  |
| Charge6 | One-hot encoded precursor charge state (1 if charge is 6, else 0) |  |
| Charge5 | One-hot encoded precursor charge state (1 if charge is 5, else 0) |  |
| Charge4 | One-hot encoded precursor charge state (1 if charge is 4, else 0) |  |
| Charge3 | One-hot encoded precursor charge state (1 if charge is 3, else 0) |  |
| Charge2 | One-hot encoded precursor charge state (1 if charge is 2, else 0) |  |
| Charge1 | One-hot encoded precursor charge state (1 if charge is 1, else 0) |  |
| ChargeN | Precursor charge state |  |
| **DeepLC retention time features** | |  |
| predicted_retention_time_best | Predicted retention time for the precursor peak (all PSMs with the same sequence, modifications, and charge) closest to the predicted retention time |  |
| observed_retention_time_best | Observed retention time for the precursor peak (all PSMs with the same sequence, modifications, and charge) closest to the predicted retention time |  |
| rt_diff_best | Difference between observed and predicted retention time for the precursor peak (all PSMs with the same sequence, modifications, and charge) closest to the predicted retention time |  |
| rt_diff | Difference between observed and predicted retention time |  |
| predicted_retention_time | Predicted retention time |  |
| observed_retention_time | Observed retention time |  |
| **MS²PIP spectrum prediction features** | |  |
| cos_iony | Cosine similarity on y-ions |  |
| cos_ionb | Cosine similarity on b-ions |  |
| cos | Cosine similarity on b- and y-ions |  |
| dotprod_iony | Dot product on y-ions |  |
| dotprod_ionb | Dot product on b-ions |  |
| dotprod | Dot product on b- and y-ions |  |
| iony_std_abs_diff | Standard deviation of the absolute differences (y-ions only) |  |
| iony_mean_abs_diff | Mean of the absolute differences (y-ions only) |  |
| iony_abs_diff_Q3 | Quantile 3 of the absolute differences (y-ions only) |  |
| iony_abs_diff_Q2 | Quantile 2 of the absolute differences (y-ions only) |  |
| iony_abs_diff_Q1 | Quantile 1 of the absolute differences (y-ions only) |  |
| iony_max_abs_diff | Maximum absolute difference (y-ions only) |  |
| iony_min_abs_diff | Minimum absolute difference (y-ions only) |  |
| ionb_std_abs_diff | Standard deviation of the absolute differences (b-ions only) |  |
| ionb_mean_abs_diff | Mean of the absolute differences (b-ions only) |  |
| ionb_abs_diff_Q3 | Quantile 3 of the absolute differences (b-ions only) |  |
| ionb_abs_diff_Q2 | Quantile 2 of the absolute differences (b-ions only) |  |
| ionb_abs_diff_Q1 | Quantile 1 of the absolute differences (b-ions only) |  |
| ionb_max_abs_diff | Maximum absolute difference (b-ions only) |  |
| ionb_min_abs_diff | Minimum absolute difference (b-ions only) |  |
| std_abs_diff | Standard deviation of the absolute differences |  |
| mean_abs_diff | Mean of the absolute differences |  |
| abs_diff_Q3 | Quantile 3 of the absolute differences |  |
| abs_diff_Q2 | Quantile 2 of the absolute differences |  |
| abs_diff_Q1 | Quantile 1 of the absolute differences |  |
| max_abs_diff | Maximum absolute difference |  |
| min_abs_diff | Minimum absolute difference |  |
| max_abs_diff_iontype | Ion type with maximum absolute difference |  |
| min_abs_diff_iontype | Ion type with minimum absolute difference |  |
| iony_mse | Mean square error on y-ions |  |
| ionb_mse | Mean square error on b-ions |  |
| spec_mse | Mean square error on b- and y-ions |  |
| iony_spearman | Spearman correlation on b- and y-ions |  |
| ionb_spearman | Spearman correlation on b- and y-ions |  |
| spec_spearman | Spearman correlation on b- and y-ions |  |
| iony_pearson | Pearson correlation coefficient on y-ions |  |
| ionb_pearson | Pearson correlation coefficient on b-ions |  |
| spec_pearson | Pearson correlation coefficient on b- and y-ions |  |
| cos_iony_norm | Cosine similarity on y-ions (log2-normalized spectrum) |  |
| cos_ionb_norm | Cosine similarity on b-ions (log2-normalized spectrum) |  |
| cos_norm | Cosine similarity on b- and y-ions (log2-normalized spectrum) |  |
| dotprod_iony_norm | Dot product on y-ions (log2-normalized spectrum) |  |
| dotprod_ionb_norm | Dot product on b-ions (log2-normalized spectrum) |  |
| dotprod_norm | Dot product on b- and y-ions (log2-normalized spectrum) |  |
| iony_std_abs_diff_norm | Standard deviation of the absolute differences (y-ions only) (log2-normalized spectrum) |  |
| iony_mean_abs_diff_norm | Mean of the absolute differences (y-ions only) (log2-normalized spectrum) |  |
| iony_abs_diff_Q3_norm | Quantile 3 of the absolute differences (y-ions only) (log2-normalized spectrum) |  |
| iony_abs_diff_Q2_norm | Quantile 2 of the absolute differences (y-ions only) (log2-normalized spectrum) |  |
| iony_abs_diff_Q1_norm | Quantile 1 of the absolute differences (y-ions only) (log2-normalized spectrum) |  |
| iony_max_abs_diff_norm | Maximum absolute difference (y-ions only) (log2-normalized spectrum) |  |
| iony_min_abs_diff_norm | Minimum absolute difference (y-ions only) (log2-normalized spectrum) |  |
| ionb_std_abs_diff_norm | Standard deviation of the absolute differences (b-ions only) (log2-normalized spectrum) |  |
| ionb_mean_abs_diff_norm | Mean of the absolute differences (b-ions only) (log2-normalized spectrum) |  |
| ionb_abs_diff_Q3_norm | Quantile 3 of the absolute differences (b-ions only) (log2-normalized spectrum) |  |
| ionb_abs_diff_Q2_norm | Quantile 2 of the absolute differences (b-ions only) (log2-normalized spectrum) |  |
| ionb_abs_diff_Q1_norm | Quantile 1 of the absolute differences (b-ions only) (log2-normalized spectrum) |  |
| ionb_max_abs_diff_norm | Maximum absolute difference (b-ions only) (log2-normalized spectrum) |  |
| ionb_min_abs_diff_norm | Minimum absolute difference (b-ions only) (log2-normalized spectrum) |  |
| std_abs_diff_norm | Standard deviation of the absolute differences (log2-normalized spectrum) |  |
| mean_abs_diff_norm | Mean of the absolute differences (log2-normalized spectrum) |  |
| abs_diff_Q3_norm | Quantile 3 of the absolute differences (log2-normalized spectrum) |  |
| abs_diff_Q2_norm | Quantile 2 of the absolute differences (log2-normalized spectrum) |  |
| abs_diff_Q1_norm | Quantile 1 of the absolute differences (log2-normalized spectrum) |  |
| max_abs_diff_norm | Ion type with maximum absolute difference (log2-normalized spectrum) |  |
| min_abs_diff_norm | Ion type with minimum absolute difference (log2-normalized spectrum) |  |
| iony_mse_norm | Mean square error on y-ions (log2-normalized spectrum) |  |
| ionb_mse_norm | Mean square error on b-ions (log2-normalized spectrum) |  |
| spec_mse_norm | Mean square error on b- and y-ions (log2-normalized spectrum) |  |
| iony_pearson_norm | Pearson correlation coefficient on y-ions (log2-normalized spectrum) |  |
| ionb_pearson_norm | Pearson correlation coefficient on b-ions (log2-normalized spectrum) |  |
| spec_pearson_norm | Pearson correlation coefficient on b- and y-ions (log2-normalized spectrum) |  |

#### Supplementary Table S4

Median Pearson correlations for each model, and evaluation data set.

| Model | Evaluation data set | Median Pearson correlation |
| --- | --- | --- |
| 2019 HCD model | Chymotrypsin | 0.756369 |
| 2019 HCD model | HLA-I | 0.805887 |
| 2019 HCD model | General proteomics | 0.910123 |
| 2019 HCD model | HLA-II | 0.791316 |
| Immuno-chymotrypsin model | Chymotrypsin | 0.898770 |
| Immuno-chymotrypsin model | HLA-I | 0.939815 |
| Immuno-chymotrypsin model | General proteomics | 0.934454 |
| Immuno-chymotrypsin model | HLA-II | 0.911232 |
| Immunopeptide model | Chymotrypsin | 0.826430 |
| Immunopeptide model | HLA-I | 0.942002 |
| Immunopeptide model | General proteomics | 0.932352 |
| Immunopeptide model | HLA-II | 0.913439 |
| Non-tryptic model | Chymotrypsin | 0.815645 |
| Non-tryptic model | HLA-I | 0.936314 |
| Non-tryptic model | General proteomics | 0.881133 |
| Non-tryptic model | HLA-II | 0.905038 |

### Supplementary figures

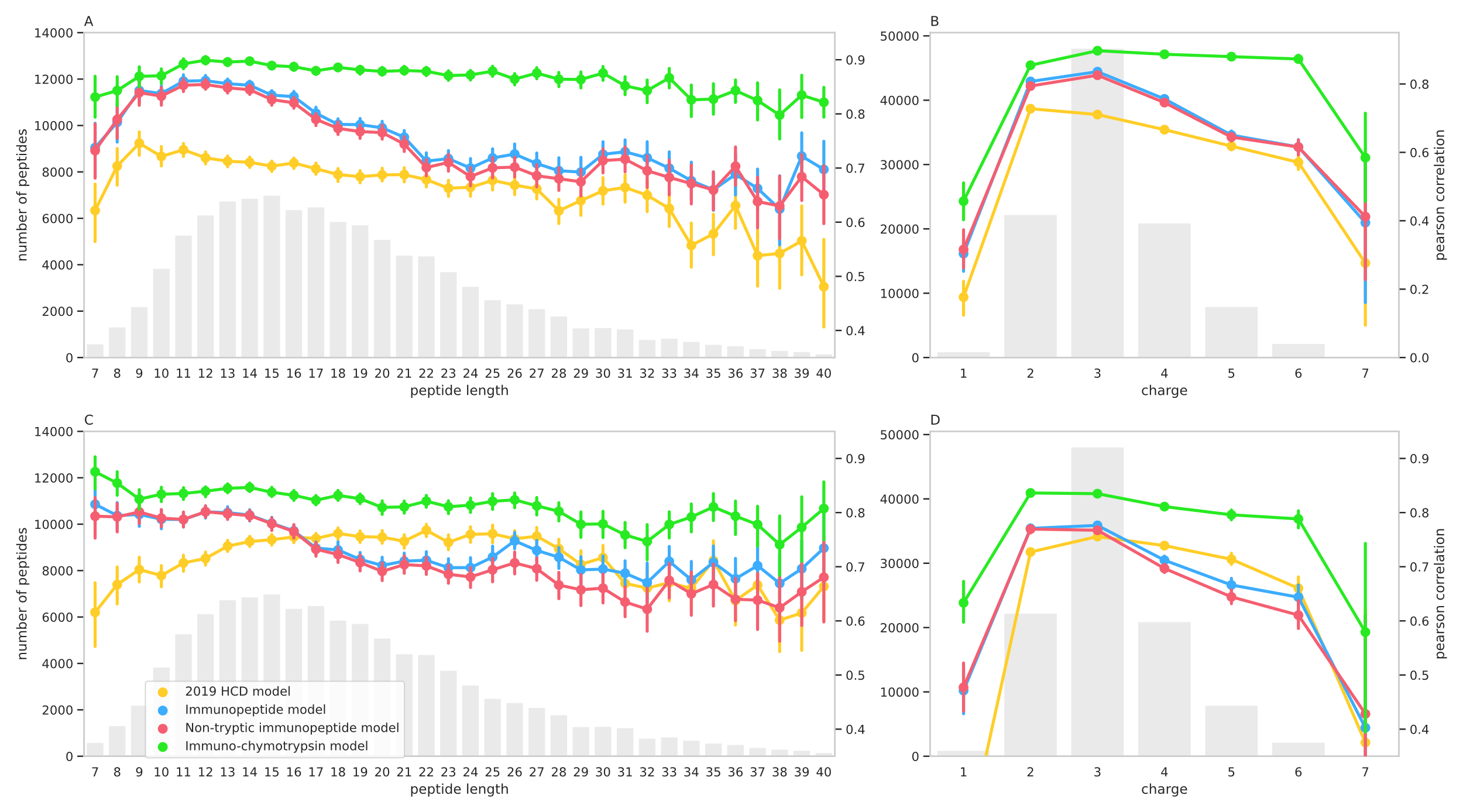

#### Supplementary Figure S1

MS²PIP model evaluation on chymotrypsin-digested data based on peptide characteristics. MS²PIP model performances on chymotrypsin-digested data split by length (A, C) and precursor charge (B, D). The connected dots indicate the median Pearson correlation for each MS²PIP y-ion model (A, B) and b-ion model (C, D). Error bars denote the Pearson correlation standard deviation within each group. The grey bar charts indicate the number of evaluation peptides within each group.

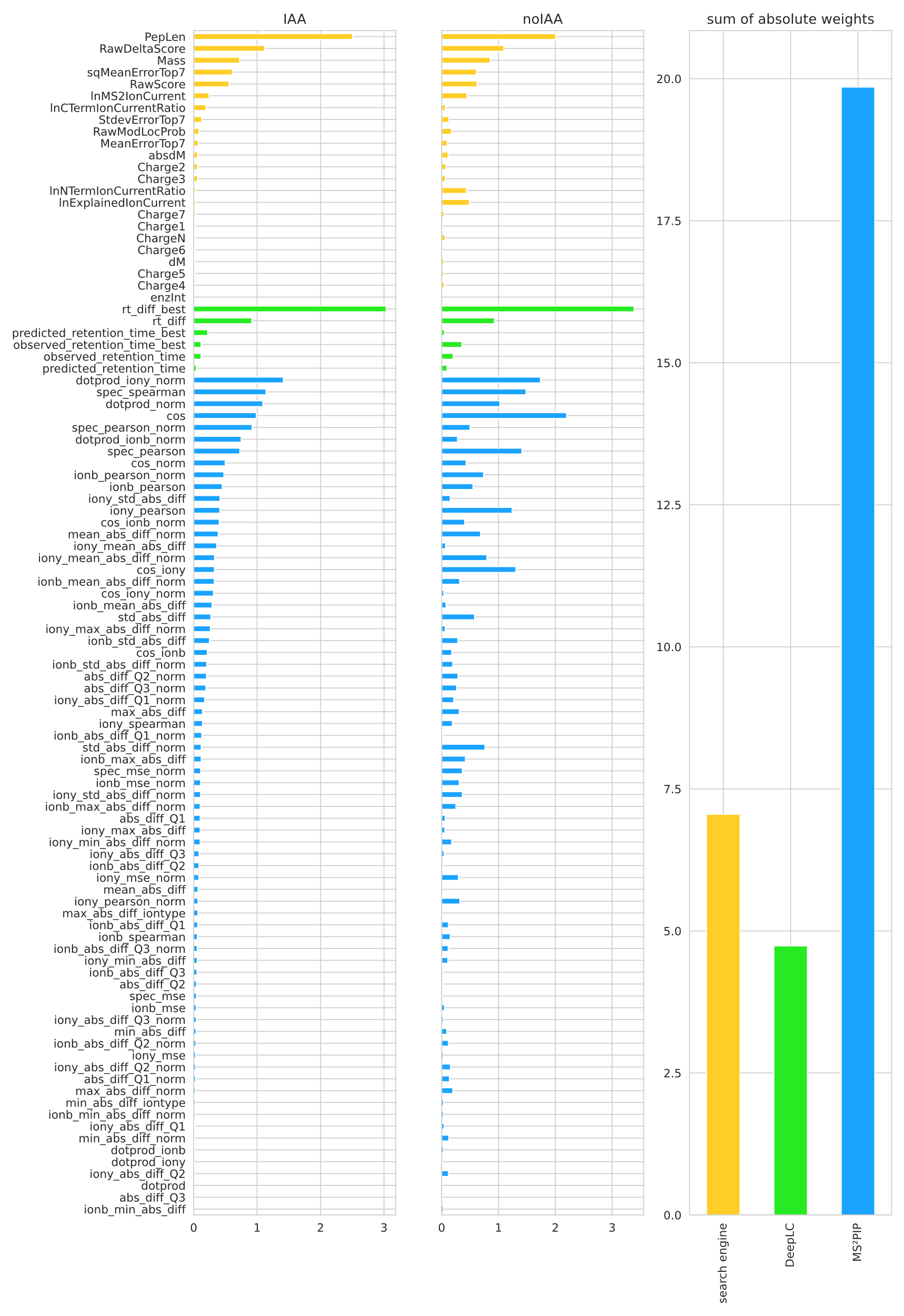

#### Supplementary Figure S2

Percolator rescoring weight analysis. Bar charts showing the mean Percolator feature weights (across the three Percolator cross validation splits) for each feature as generated by MS²Rescore, split by alkylated samples (IAA) on the left and non-alkylated samples (noIAA) in the middle. Blue bars denote MS²PIP based-features, green bars denote DeepLC-based features and yellow bars denote search engine derived features. The bar chart on the right shows the combined weight of each category, calculated by summing the respective absolute weights across both samples.

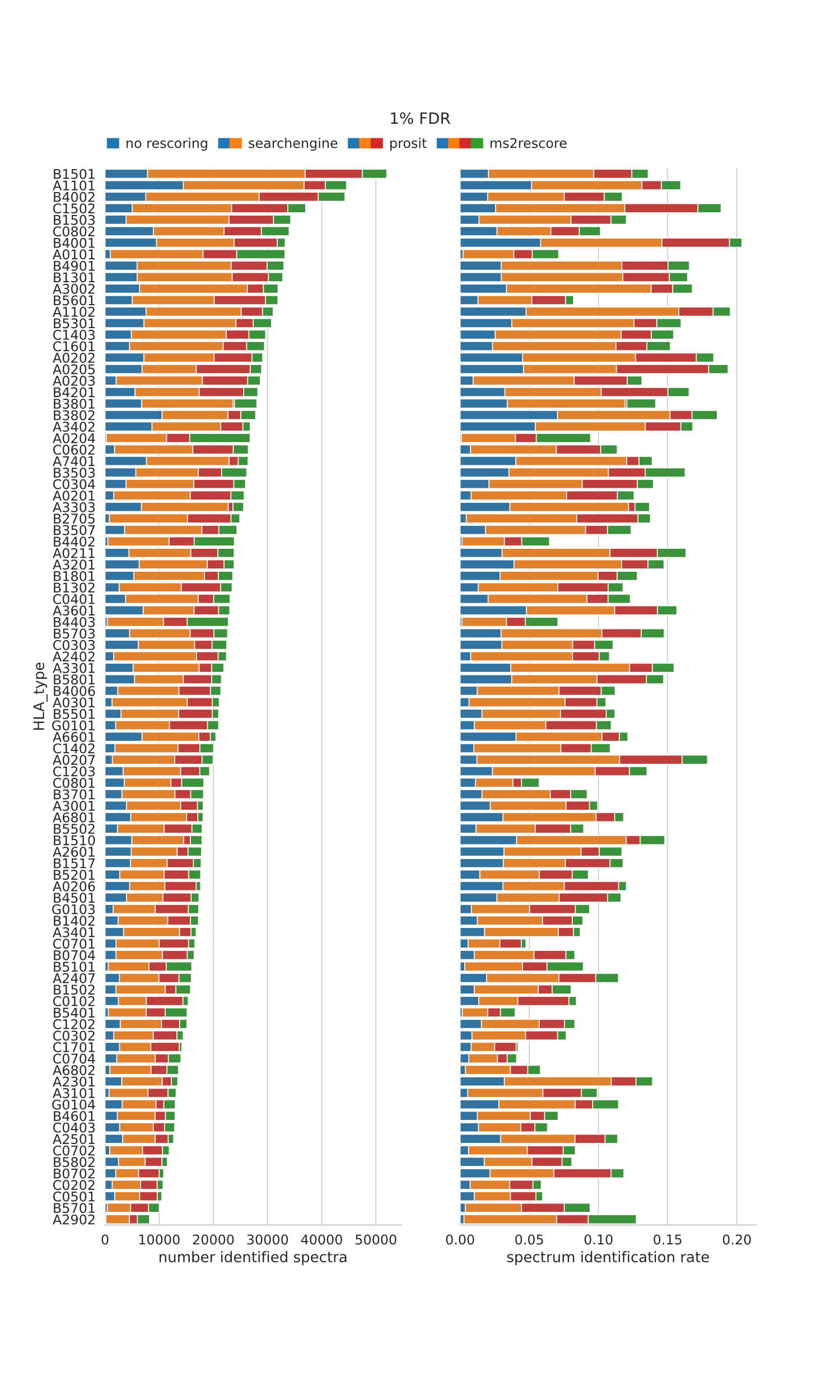

#### Supplementary Figure S3

Number of identifications and identification rate based on HLA pattern for several rescoring methods for 1%FDR.The numbers of identified spectra are shown on the left with the corresponding identification rate shown on the right.

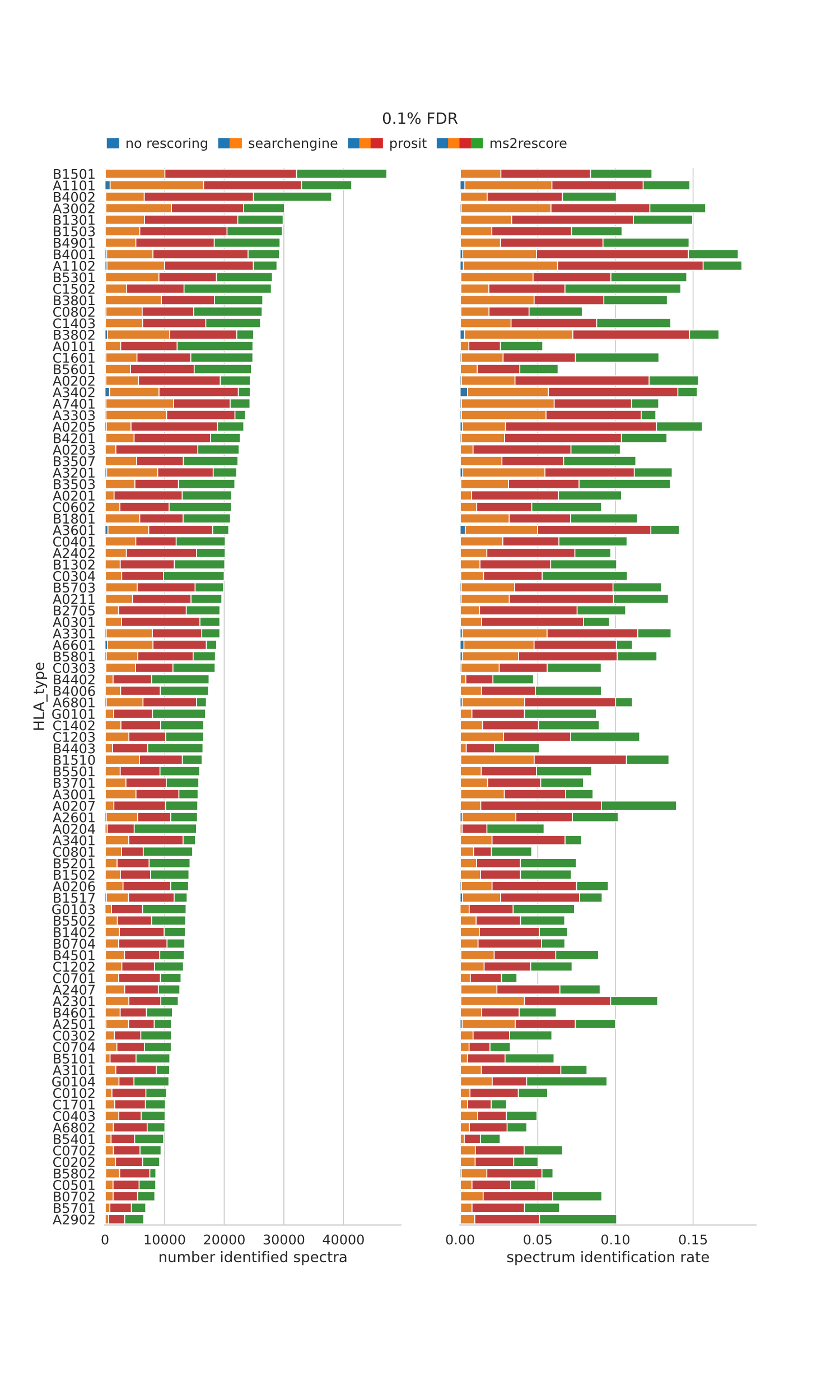

#### Supplementary Figure S4

Number of identifications and identification rate based on HLA pattern for several rescoring methods for 0.1%FDR.The numbers of identified spectra are shown on the left with the corresponding identification rate shown on the right.

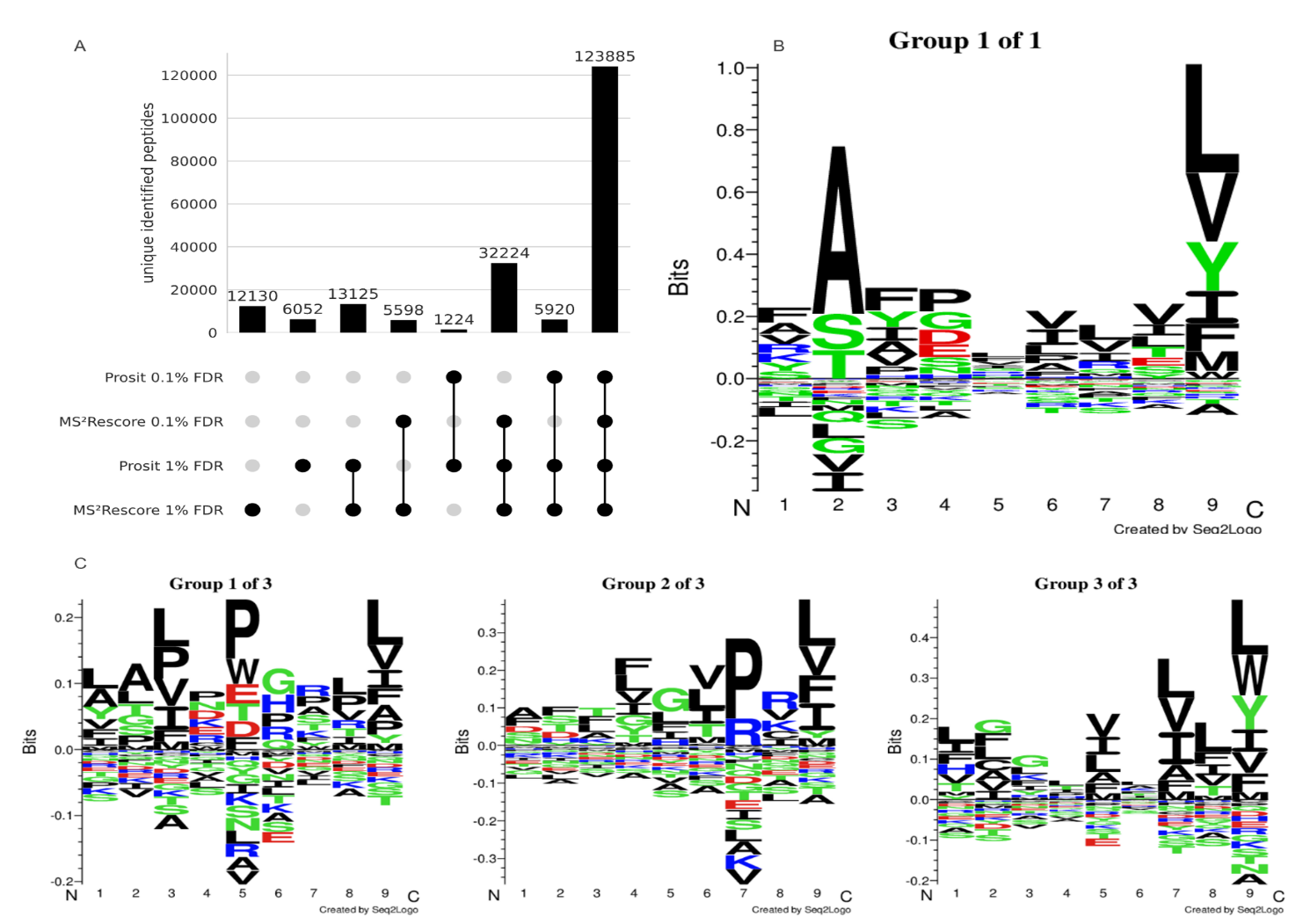

#### Supplementary Figure S5

Comparison of Prosit and MS²Rescore for 1% and 0.1%. Upset plot showing the unique identified spectra of MS²Rescore and Prosit.

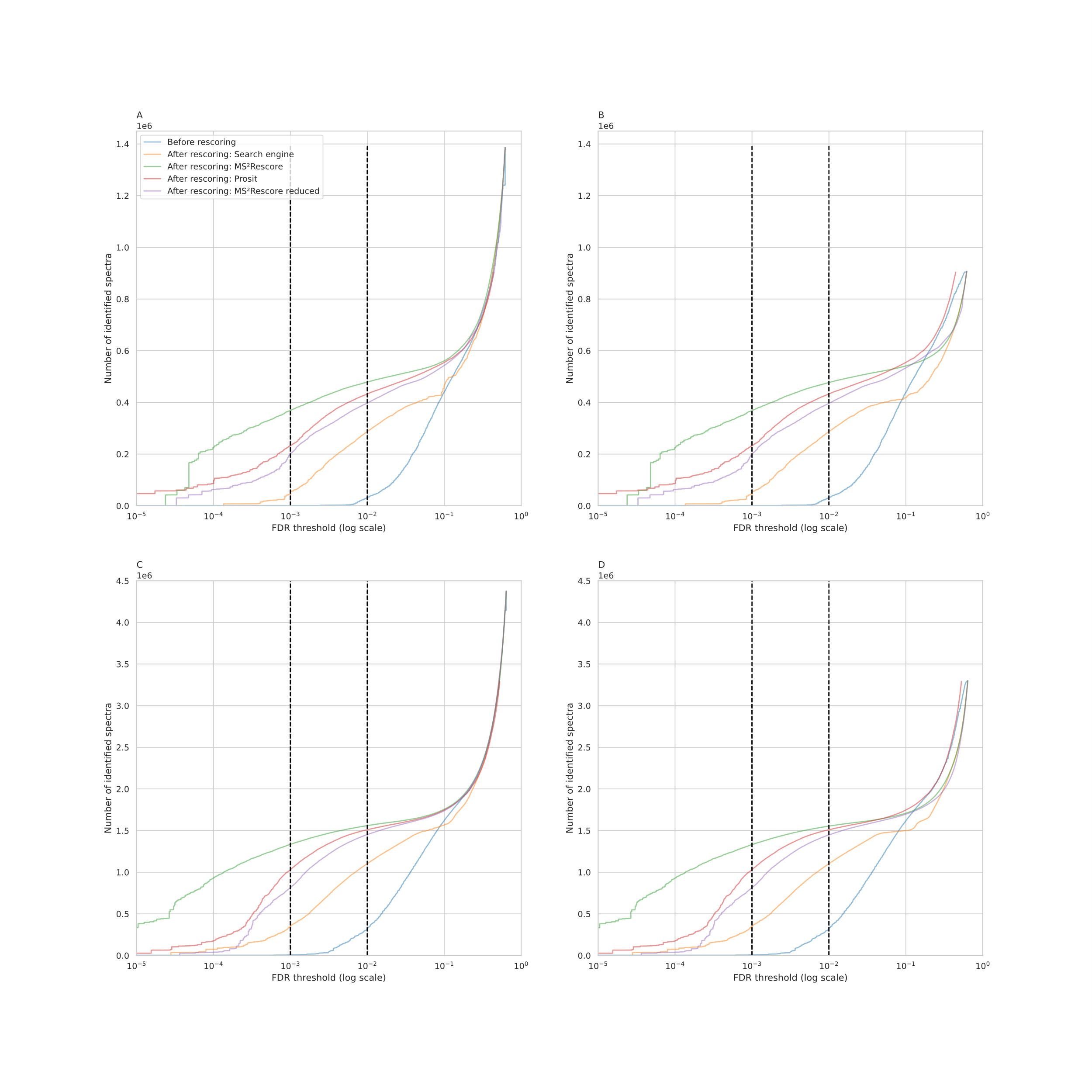

#### Supplementary Figure S6

Comparison of the total number of identified spectra for different rescoring methods at varying FDR thresholds in log10 scale. A, B, C, D shown rescoring methods used are: Raw MaxQuant (Andromeda) scores without rescoring (Before rescoring, blue), rescoring with only search engine feature as Percolator would do (After rescoring: Search engine, orange), full MS²Rescore rescoring with search engine-, DeepLC-, and MS²PIP-features (After rescoring: MS²Rescore, green), rescoring with Prosit as performed by the original authors (After rescoring: Prosit, red), and MS²Rescore without DeepLC retention time features and most search engine features, emulating the Prosit feature set (After rescoring: MS²Rescore reduced, purple). The dotted black lines denote the 0.1% false discovery rate threshold (left) and 1% false discovery rate threshold (right). Figures A and C show results for the non-alkylated samples (noIAA), figures B and D results for the alkylated samples (IAA). Figures on the left show results including all identifications compatible with MS²Rescore, figures on the right show results filtered down to only including identifications compatible with Prosit, which is a subset of the former.

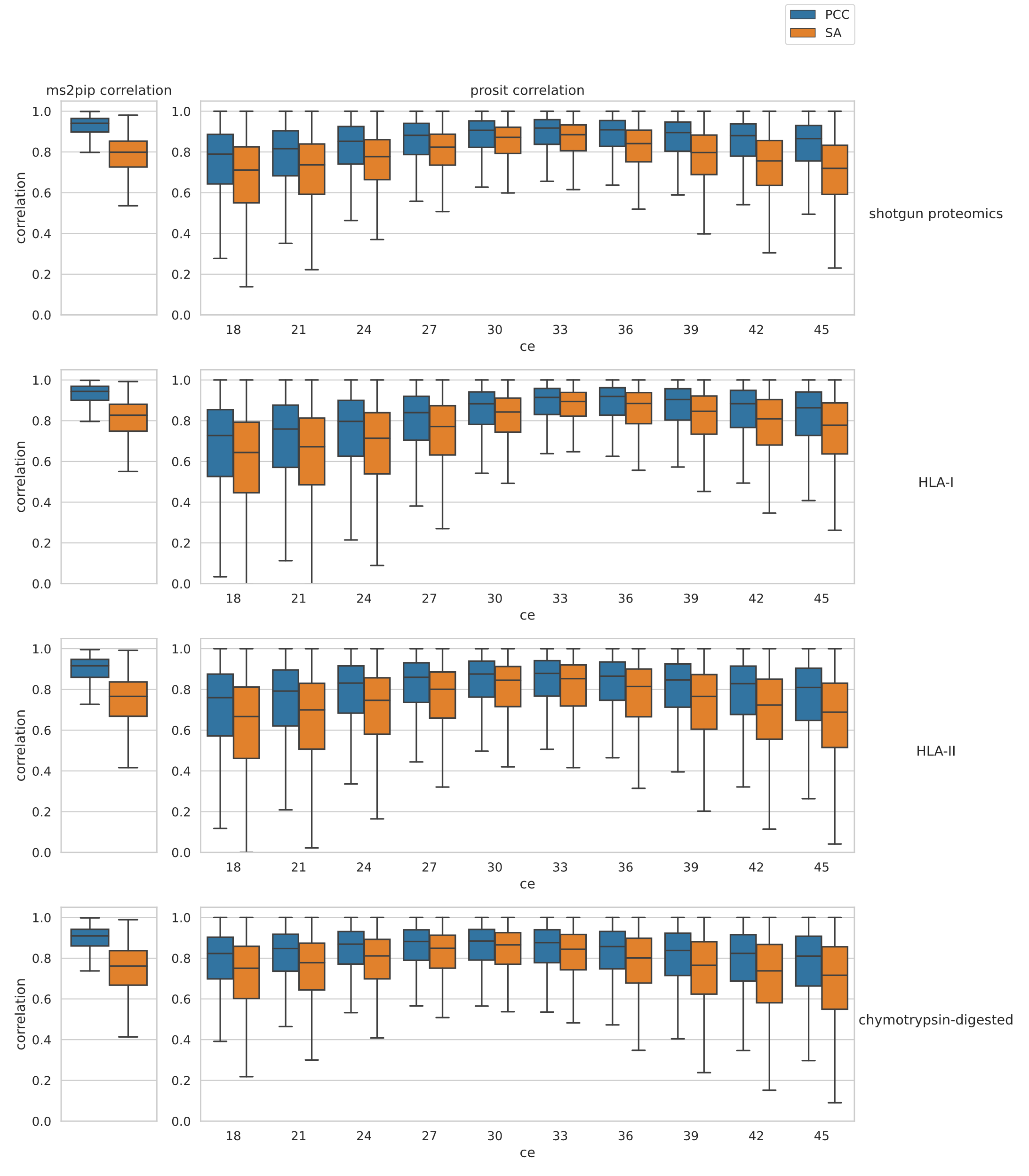

#### Supplementary Figure S7

Comparison of peak intensity prediction correlations between MS²PIP and Prosit for the different evaluation data sets that were also used in Figure 1. The boxplots show the Pearson correlation coefficient (PCC) and spectral angle (SA) for different evaluation sets per PSM for MS²PIP (left) and Prosit (right). For Prosit, the predictions are shown for different collision energy settings in the model. Intensities for +2 and +3 fragment ions were left out of the comparison, as MS²PIP only predicts intensity for singly charged fragment ions.

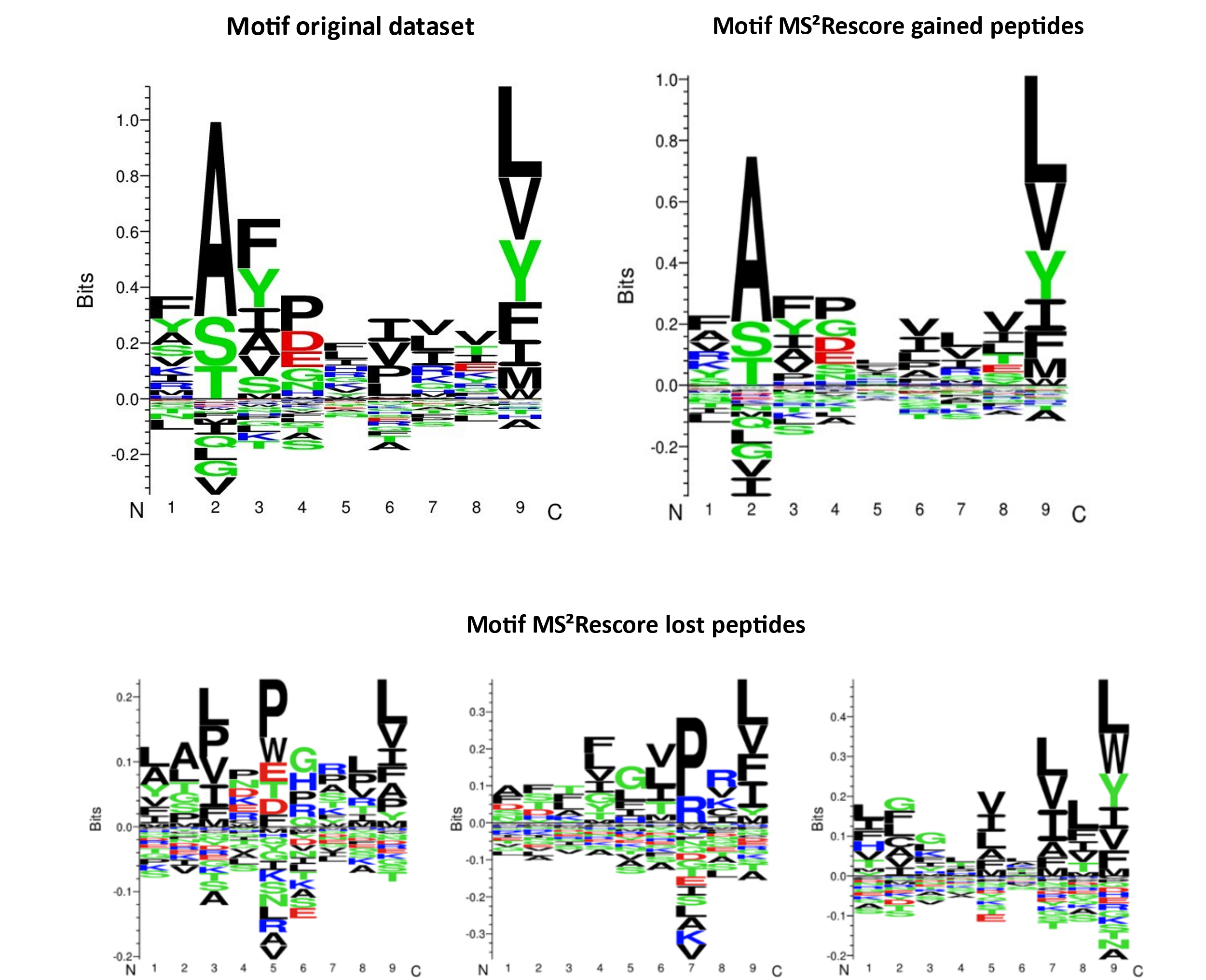

#### Supplementary Figure S8

Comparison of Prosit and MS²Rescore for 1% and 0.1% FDR and motif analysis of MS²Rescore identifications. Sequence motif(s) shown for the C*12:03 HLA type calculated for the original dataset (Sarkizova & Klaeger et al) (A), the gained (B) and lost (C) sequences identified by MS²Rescore in relation to search engine rescoring below the 1% FDR threshold. All figures are created by Seq2Logo through GibbsCluster (v2.0).

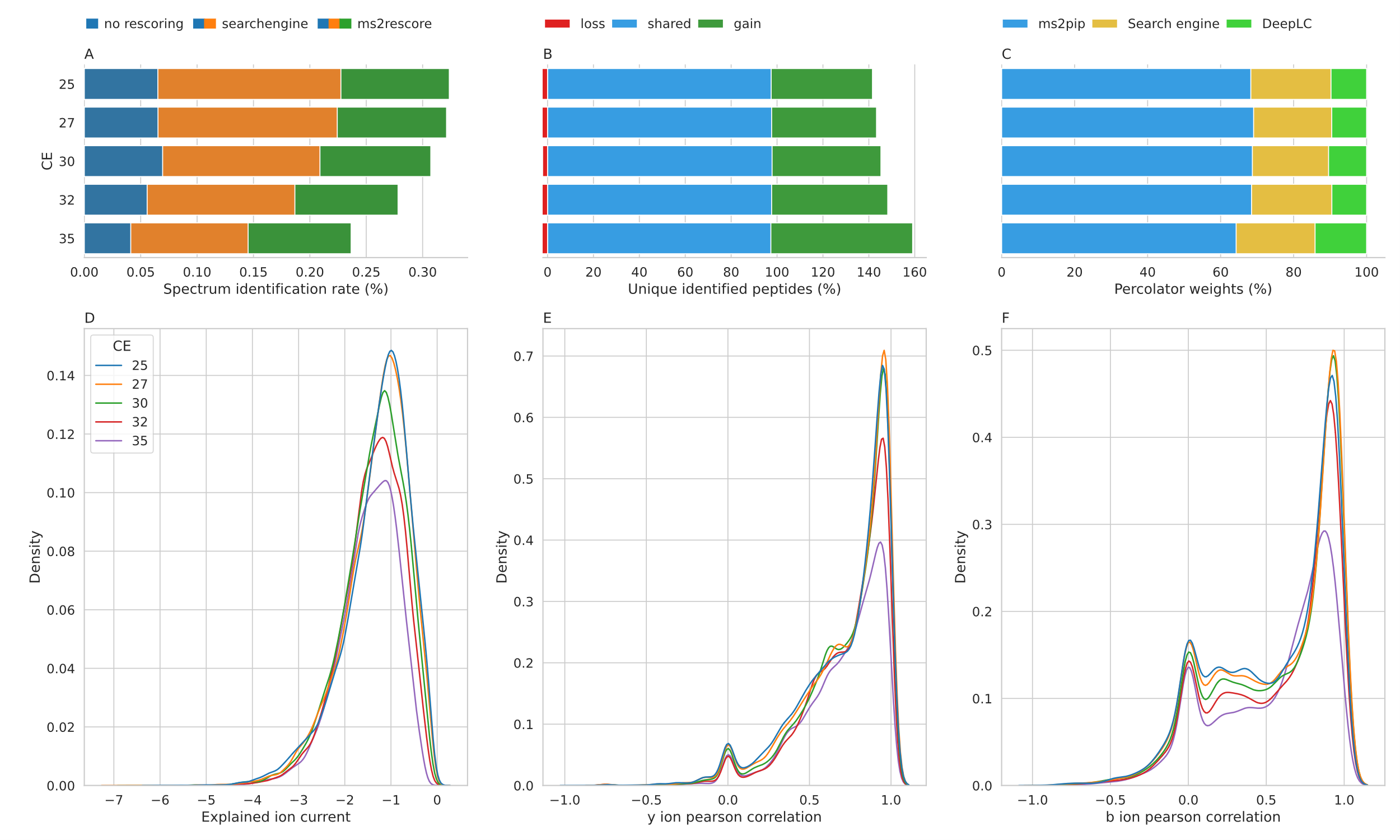

#### Supplementary Figure S9

MS²Rescore analysis for various collision energy settings. Bar charts showing the spectrum identification rate for no rescoring, search engine rescoring and rescoring with all features from MS²Rescore for various collision energies (A), showing the shared (blue), gained (green) and lost (red) number of unique (by sequence) identified immunopeptides in relation to rescoring with only search engine features for the 1% FDR threshold (B), and showing the relative weight of the Percolator features for each feature set for various collision energies (C). Density plots showing the explained ion current, i.e. how much intensity comes from the peptide fragmentation, for various collision energies (D), showing the y (E) and b (F) ions pearson correlation between observed and predicted spectra separated per collision energy value. (note all zero values are caused by the fact that either the observed or predicted intensities for a given ion type are all zero).

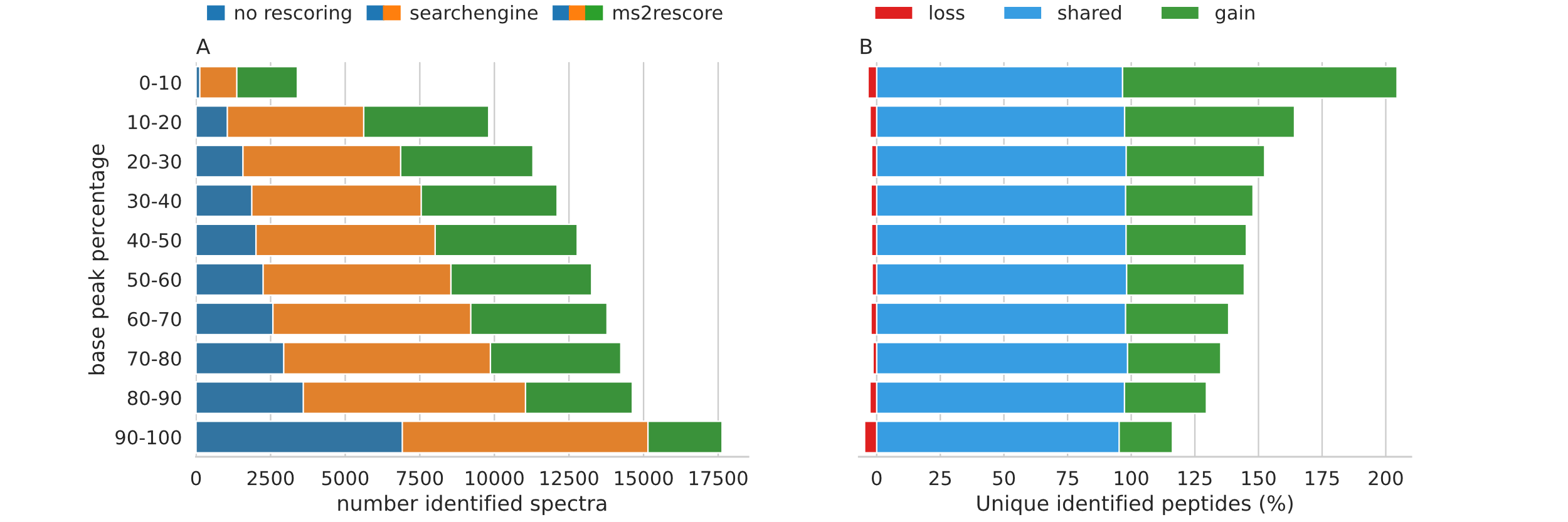

#### Supplementary Figure S10

Evaluation of MS²Rescore across peptide abundances. Bar charts split by MS1 peak intensity discretized in 10 deciles showing (A) the number of identified spectra for each feature set and (B) the number of shared, gained, and lost unique identified immunopeptides in terms of sequence, compared to rescoring with only search engine features for the 1% FDR threshold.

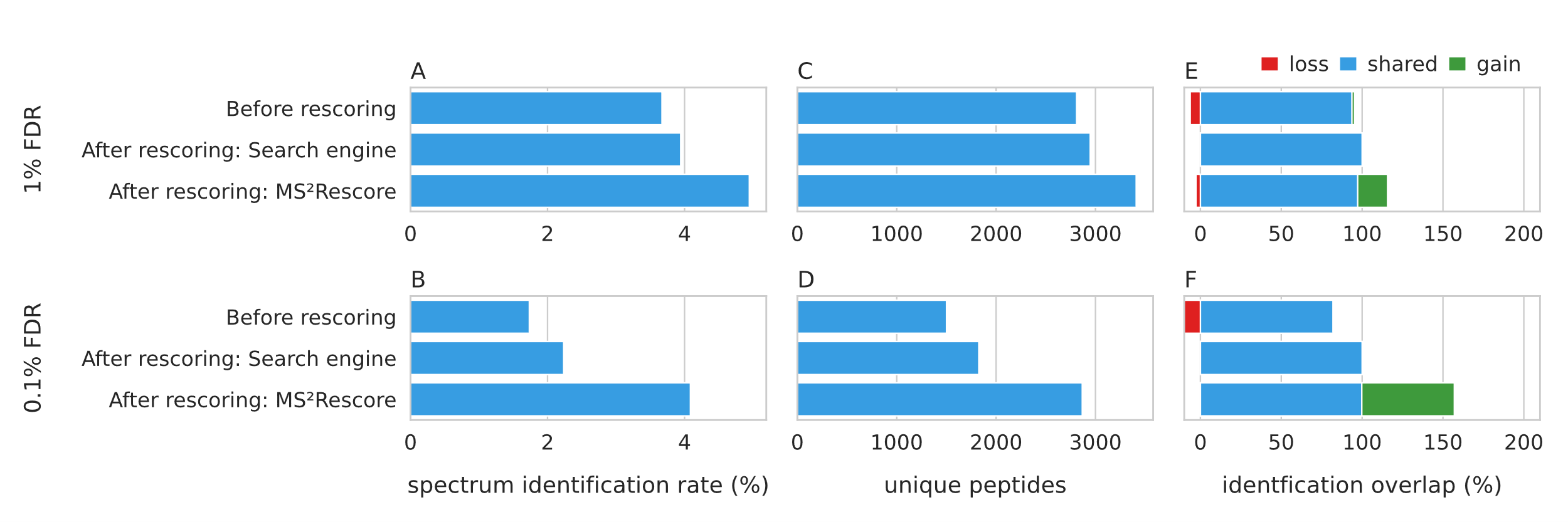

#### Supplementary Figure S11

Percentage of identified spectra and unique identified peptides using different rescoring methods for HLA class II peptides (PXD015408) processed with the PEAKS DB search engine. Bar charts showing the spectrum identification rate out of 198.669 spectra (A, B), showing total number of unique identified peptides in terms of sequence for each rescoring method (C, D) and showing the Bar chart showing the shared (blue), gained (green) and lost (red) number of unique (by sequence) identified immunopeptides (E, F) in relation to rescoring with only search engine features. All results are shown for the 1% FDR (A, C, E) and 0.1% FDR (B, D, F) thresholds.
